# *Pseudomonas aeruginosa* faces a fitness trade-off between mucosal colonization and antibiotic tolerance during airway infections

**DOI:** 10.1101/2024.09.09.611974

**Authors:** Lucas A. Meirelles, Evangelia Vayena, Auriane Debache, Eric Schmidt, Tamara Rossy, Tania Distler, Vassily Hatzimanikatis, Alexandre Persat

**Author notes:** Department of Mechanical Engineering, Massachusetts Institute of Technology, Cambridge, MA, USA.

## Abstract

*Pseudomonas aeruginosa* causes antibiotic-resilient acute and chronic pneumonia, but the mechanisms by which it adapts to the airway environment are poorly understood. Here, we investigated *P. aeruginosa* pathoadaptive mechanisms in tissue-engineered human airway organoids. Using transposon sequencing *in situ,* we decoded how *P. aeruginosa* survives on the mucosal surface during antibiotic treatment. Biofilm formation emerged as a major driver of *P. aeruginosa* colonization. Mutants that extensively produce biofilms on mucus show limited exploratory behavior, which limits nutrient access, slowing down their growth. Conversely, biofilm-dwelling *P. aeruginosa* better tolerate antibiotics via biophysical mechanisms. Finally, biofilms can shelter less-tolerant but more cytotoxic strains, thereby contributing to genotypic heterogeneity. *P. aeruginosa* must therefore adapt to conflicting physical and biological selective pressures to initiate chronic infections.

## Introduction

The opportunistic pathogen *Pseudomonas aeruginosa* is a major contributor to the current antimicrobial resistance crisis. *P. aeruginosa* is inherently tolerant to antimicrobials, and multidrug and pandrug-resistant strains are on the rise ^1,2^. As a result, new therapeutic strategies to fight off *P. aeruginosa* infection are now a top priority ^3,4^. *P. aeruginosa* causes severe acute and chronic pneumonia in immunocompromised patients. Individuals with obstructive lung diseases such as cystic fibrosis (CF) and chronic obstructive pulmonary disease (COPD) are particularly at risk for developing persistent infections with high mortality rates ^5,6^. Despite this dire situation, we still know little about the mechanisms used by *P. aeruginosa* to adapt to the lung environment, thereby impeding the rational development of novel anti-infectives.

Throughout infection, host factors and antibiotics exert strong selective pressures on microbial populations, which result in evolutionary trajectories that ultimately select for recalcitrance to antibiotic treatment and chronicity ^7^. Early in this process, *P. aeruginosa* phenotypically and genotypically transitions from an acute state characterized by a planktonic lifestyle with high cytotoxicity to a chronic state commonly associated with the formation of multicellular aggregates called biofilms ^7^. However, acute strains can re-emerge during chronic infections, frequently resulting in tissue damage and a decline in lung function ^7–9^. The selective forces and fitness costs controlling these transitions and cycles are still poorly understood ^10^.

To form biofilms, bacteria embed themselves in a self-secreted matrix of extracellular polymeric substances (EPS). Psl and Pel polysaccharides are the main components of the *P. aeruginosa* EPS matrix, both of which are regulated by the second messenger cyclic-di-GMP ^11–13^. Biofilm-dwelling cells can be orders of magnitude more tolerant to common antibiotics than planktonic cells ^14–17^, which increases *P. aeruginosa*’s likelihood of evolving drug resistance ^15,17,18^. Clinical isolates from chronic *P. aeruginosa* pneumonia are dominated by mutants hyper-secreting EPS matrix ^7,19–21^, often due to defects in cdGMP signaling. However, the mechanisms leading to the selection of these biofilm-forming isolates are poorly understood. Our limited understanding arises from the fact that this process has mainly been investigated in the context of animal models that do not replicate human physiology and limit mechanistic investigations at high temporal and spatial resolutions ^22,23^. Conversely, most mechanistic studies of *P. aeruginosa* physiology in laboratory conditions overlook biological and physical factors from the host. To initiate infection, *P. aeruginosa* first encounters mucus, a hydrogel substance lining the airway epithelium. Airway mucus is composed of cross-linked gel-forming mucins MUC5B and MUC5AC, which hydrate upon exocytosis ^6,24^. The mucus layer normally protects host cells from particles and microbes in a process known as mucociliary clearance ^6,25^. Individuals with CF and COPD have abnormally viscous mucus, leading to accumulation in the respiratory tract that is thought to favor *P. aeruginosa* colonization ^6,7^. In addition to being a food source, mucus constitutes a physical substrate for *P. aeruginosa* attachment close to epithelial cells, providing a favorable environment for *P. aeruginosa* colonization and chronic infections ^26^. However, the mechanisms by which *P. aeruginosa* thrives at the mucosal surface remain unresolved due to our limited ability to emulate the airway environment in the lab. Natural or synthetic sputum has been used to investigate *P. aeruginosa*’s biochemical adaptations to the airway ^27–30^. These have revealed important factors of infection but overlook local physical and spatial aspects of the mucosal surface. To bridge the gap between test tubes and clinical studies, we recently developed AirGels: a tissue-engineered 3D human lung organoid model that replicates biophysical features of the airway environment and that is suitable for high-resolution live microscopy ^26^.

Here, we elucidate how *P. aeruginosa* adapts to the mucosal surface to transition from acute to chronic states. We leverage a multidisciplinary approach combining AirGels with functional genomics, metabolic modeling, and live tracking of infections to track bacterial fitness at the mucosal surface. We identified that *P. aeruginosa* faces metabolic and biophysical fitness trade-offs as it forms biofilms under the pressure of mucosal growth and antibiotic treatment.

## Results

### Functional genomics identifies how P. aeruginosa adapts to mucosal environment

To identify how *P. aeruginosa* thrives at the mucosal surface, we performed transposon-insertion sequencing (Tn-seq) on populations growing at the surface of human primary bronchial epithelial cells (HBE cells) from a healthy donor ^31^. These human-derived primary tissues possess *in vivo*-like cell types and secrete mucus at the air-liquid interface ^26,32,33^. We first considered infection times where *P. aeruginosa* predominantly grows and colonizes mucus without affecting epithelial cell viability (Fig. 1A, Fig. S1A). We thus grew a *P. aeruginosa* Tn5-based transposon library at the mucosal surface of HBE cultures (Fig. 1B, Fig. S1B). We then sequenced the populations of transposon mutants to identify selected mutants and the pressures imposed onto them at the mucosal surface. Controls include the original library, the inoculum used for the infections, and liquid culture grown for the same duration as the infection (Fig. S1B). Tn-seq identified 620 genes whose mutation impacted *P. aeruginosa*’s fitness at the mucosal surface compared to the liquid culture reference (Table S1). This relatively high number suggests that at the mucosal surface, *P. aeruginosa* experiences selective pressures that differ vastly from typical laboratory conditions (Fig. 1C, Table S1). As expected, genes coding for metabolism were well-represented (Fig. 1C). Also, we found a surprisingly strong selection against mutants in genes coding for biofilm formation and cdGMP signaling (Fig. 1C, Table S1).

**Figure 1.**
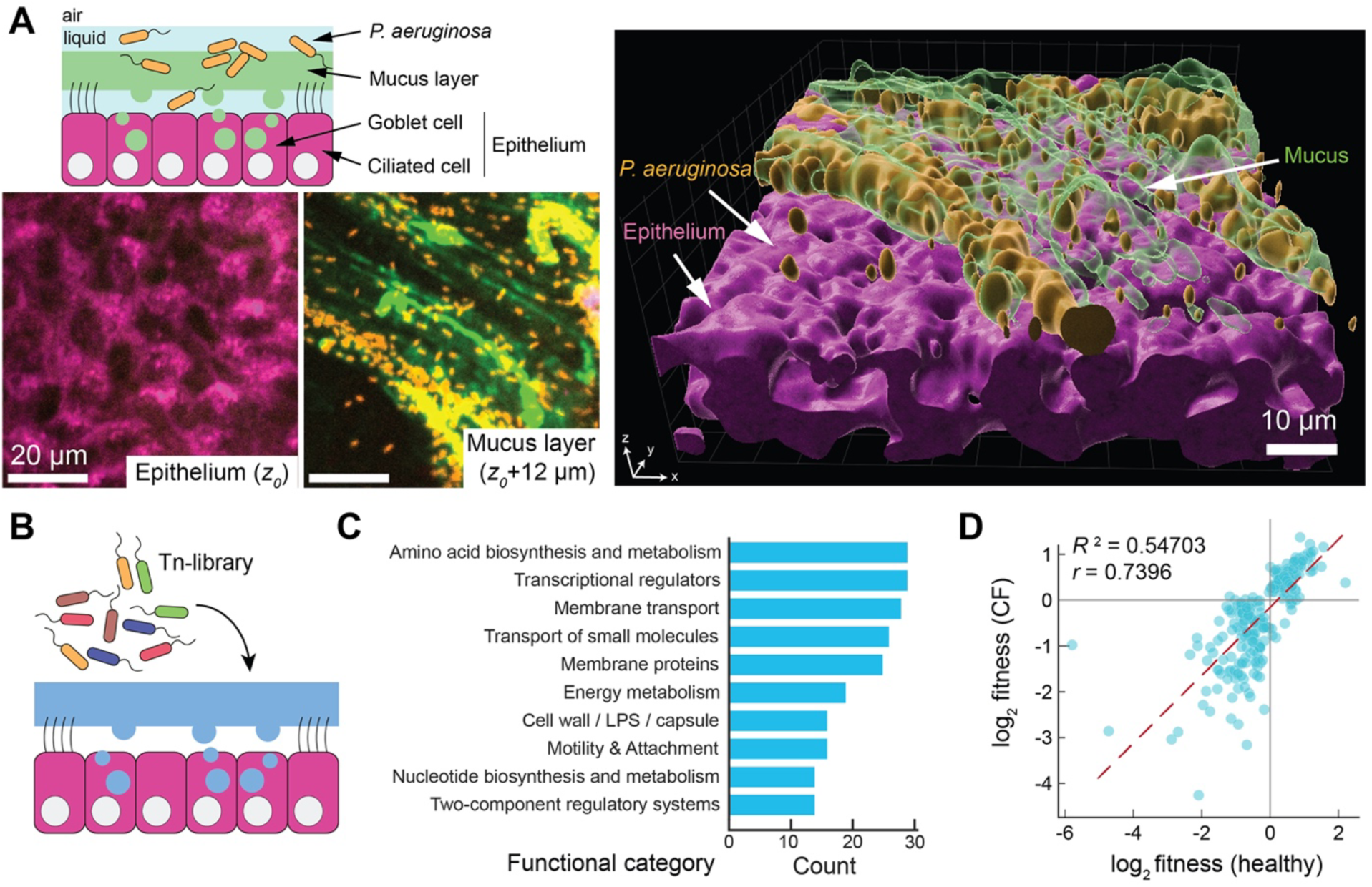
Functional genomics identifies how *P. aeruginosa* adapts to the mucosal surface. **A.** Human primary bronchial epithelial (HBE) cells differentiated at the air-liquid interface as a model for *P. aeruginosa* infections. Simplified model (top left) with confocal image (bottom left) and 3D rendering (right) of *P. aeruginosa* growing on the mucus layer without epithelial cell damage. **B.** Simplified representation of the Tn-seq experiment with the transposon library growing on the mucosal layer. **C.** Functional classification of genes found in the Tn-seq (PseudoCAP), not including genes classified as hypotheticals or coding for non-determined “putative enzymes”. The dataset used here comprises Tn-seq performed on non-CF HBE in comparison to the inoculum reference. **D.** Correlation between *P. aeruginosa* mutant fitness growing on CF and non-CF HBE cultures. Each data point represents a gene found in the Tn-seq. Only genes that passed the significance cutoff (p < 0.05) in at least one of the conditions are shown. Dashed line: linear fit for the data. *r*: Pearson correlation coefficient. *R*^2^: coefficient of determination.

Given the different evolutionary trajectories observed in the clinical setting ^10,34^, we wondered whether non-CF and CF mucosal surfaces impose distinct selective pressures on *P. aeruginosa* mutants. We, therefore, performed a similar Tn-seq analysis on HBE cultures produced from the primary cells of a CF donor. Out of the 252 mutants that passed the significance threshold (Fig. S1B), 238 were selected for or against in the same manner in both cultures (Fig. 1D, Table S2). Overall, fitness changes caused by transposon insertions correlated very strongly between CF and non-CF HBE cultures (Fig. 1D), suggesting they impose similar selective pressures on *P. aeruginosa*. In particular, biofilm production was selected against in the CF HBE cultures despite their clinical dominance during chronic infections.

### P. aeruginosa adopts metabolic independence at the mucosal surface

We used the Tn-seq data as a basis for resolving how *P. aeruginosa* thrives to the complex physicochemical environment of the airway. Due to their prevalence, we particularly focused on investigating the mechanisms by which metabolic traits and biofilm formation influence *P. aeruginosa* fitness at the mucosal surface. Mutations in genes involved in amino-acid and nucleotide biosynthesis resulted in strong fitness defects, indicating that *P. aeruginosa* is starving for those nutrients. In addition, mutants that lost functional gene products involved in glucose and lactate utilization were outcompeted. These included *edd*, participating in the Entner-Doudoroff (ED) pathway, and *lldP*, a lactate permease ^35,36^. This suggests that *P. aeruginosa* may preferentially utilize glucose and lactate as carbon sources when colonizing the mucosa (Fig. 2A). Mutants that lost the ability to make the siderophore pyoverdine *pvdE* transporter were also selected against, showing that *P. aeruginosa* starves for iron under these conditions (Fig. 2A) ^37^. We validated the contributions of *edd*, *lldP,* and *pvdE* by directly measuring the growth of corresponding in-frame clean deletion mutants with colony forming units (CFU). While these mutant strains grew as well as WT in liquid cultures, they showed lower fitness at the mucosal surface, confirming this was not a result of generic growth defects (Fig. 2B). Thus, during mucosal colonization, *P. aeruginosa* must synthesize essential compounds that cannot be easily acquired from its host, a process known as metabolic independence^38^.

**Figure 2.**
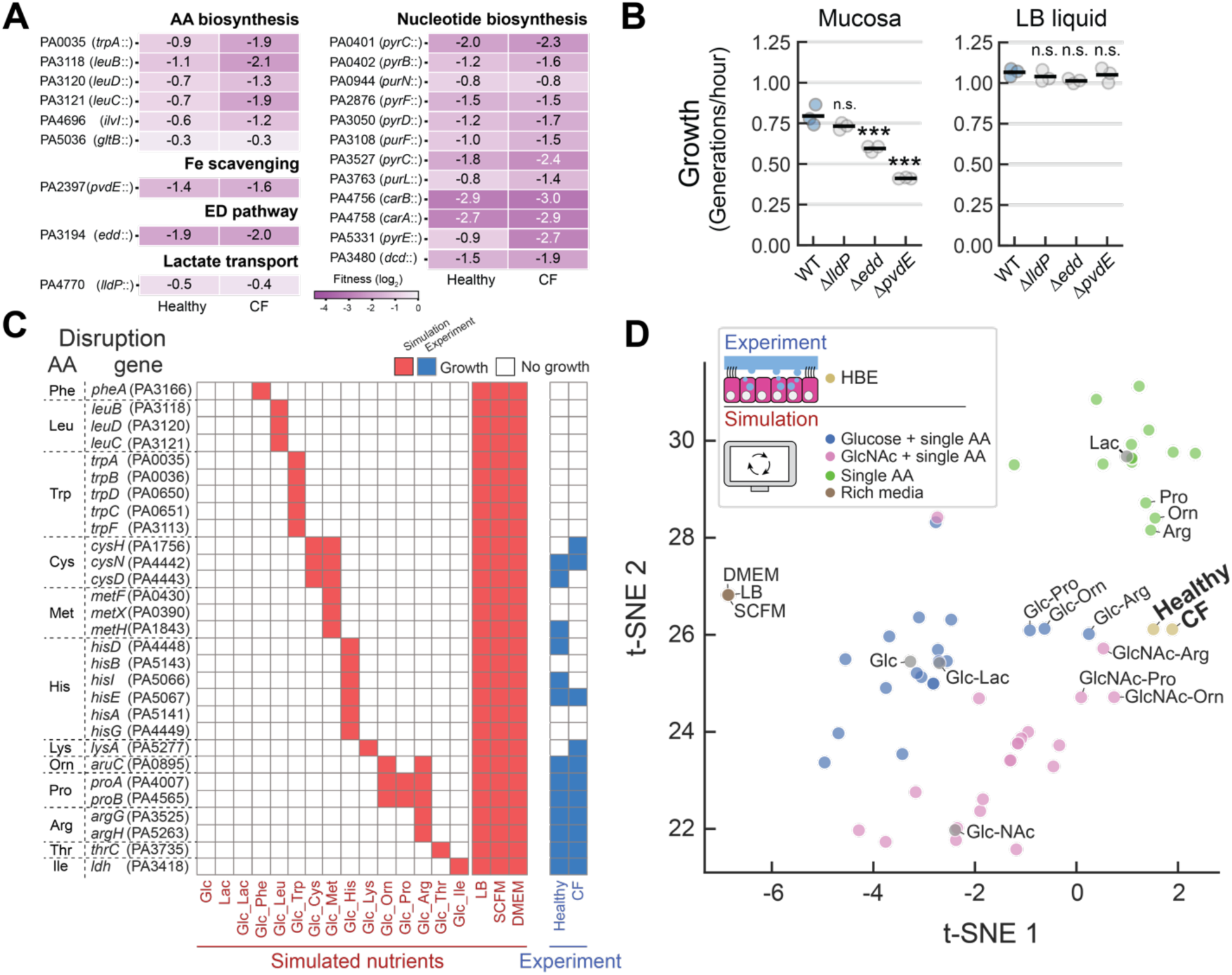
*P. aeruginosa* adopts metabolic independence at the mucosal surface. **A.** Fitness defects of transposon insertions in representative metabolic genes. **B.** Fitness of chromosomal clean deletion mutants in metabolic genes at the mucosa (left) and in LB liquid (right) by colony forming units (CFU). Each data point represents an independent biological replicate (*n* = 3); the horizontal black lines mark their mean. **C.** Metabolic network modeling simulates the effect of disrupting genes (left) during growth in simulated media conditions (bottom). The red square indicates the mutant grew in the simulation. Comparison with growth at the mucosal surface from experimental Tn-seq data is in blue. **D.** t-distributed Stochastic Neighbor Embedding (t-SNE) visualization of the Tn-seq experimental dataset and the metabolic simulations. Each data point represents one condition (simulated or experimental). Statistics: panel B, 1-way ANOVA with Tukey HSD multiple comparison test, with asterisks showing significant differences relative to WT (*** *p* < 0.001, n.s. *p* > 0.05).

To identify *P. aeruginosa*’s metabolic genes needed for fitness, we turned to metabolic network modeling. This approach gives us hints of what the nutritional environment is like at the mucosa. We performed *in silico* gene essentiality analysis using a genome-scale metabolic model of *P. aeruginosa* ^39^. We computationally analyzed gene essentiality in 61 minimal and rich media compositions spanning a wide range of carbon sources and amino acid compositions. We then compared *in silico* fitness to the one experimentally measured by Tn-seq (Table S2). We observed that while the biosynthesis of most amino acids was essential for *P. aeruginosa* to grow on the mucosal surface, mutations in genes involved in the biosynthesis of ornithine, proline, and arginine did not produce fitness defects (Fig. 2C), indicating those nutrients could be available at the surface of HBE cultures (Fig. 2C).

To further illuminate nutrient availability at the mucosal surface, we performed dimensionality reduction and clustering of both simulated and experimental data. This analysis shows that *P. aeruginosa* metabolism in HBE cultures differs from liquid culture conditions, including those aiming at replicating the composition of sputum (Fig. 2D). Simulations with a single carbon source clustered far from the Tn-seq dataset (Fig. 2D). Simulated growth in minimal media supplemented with glucose, N-Acetylglucosamine (Glc-NAc) and three specific amino acids (ornithine, proline, arginine) clustered with our data most closely. In addition to the utilization of glucose and/or lactate, the potential utilization of Glc-NAc suggests that *P. aeruginosa* may feed from glycan components of mucins that compose the mucus layer.

### Biofilm formation incurs a fitness penalty during mucosal colonization

While *P. aeruginosa* biofilms are often associated with chronic lung colonization ^40^, mutations that inhibit their formation were enriched in our Tn-seq data (Fig. 3A). In particular, loss-of-function mutants in EPS matrix component genes such as the *psl* operon improved fitness in HBE cultures (Fig. 3A). Mutations affecting biofilm regulatory components were also differentially selected at the mucosal surface. Most notably, genes regulating intracellular levels of cdGMP underwent strong selection. Mutants with low levels of cdGMP were fitter on mucus relative to liquid culture (Fig. 3A and Table S2) ^41,42^. Conversely, mutants with elevated cdGMP ^13,43–45^, such as Tn-insertion in the *wspF* gene, showed severe fitness defects (Fig. 3A). Competing WT with an in-frame clean deletion mutant Δ*wspF* confirmed this prediction: WT rapidly overtook Δ*wspF* when growing at the mucosa (Fig. 3B, left), while both strains grew at the same rate in liquid LB cultures (Fig. 3B, right).

**Figure 3.**
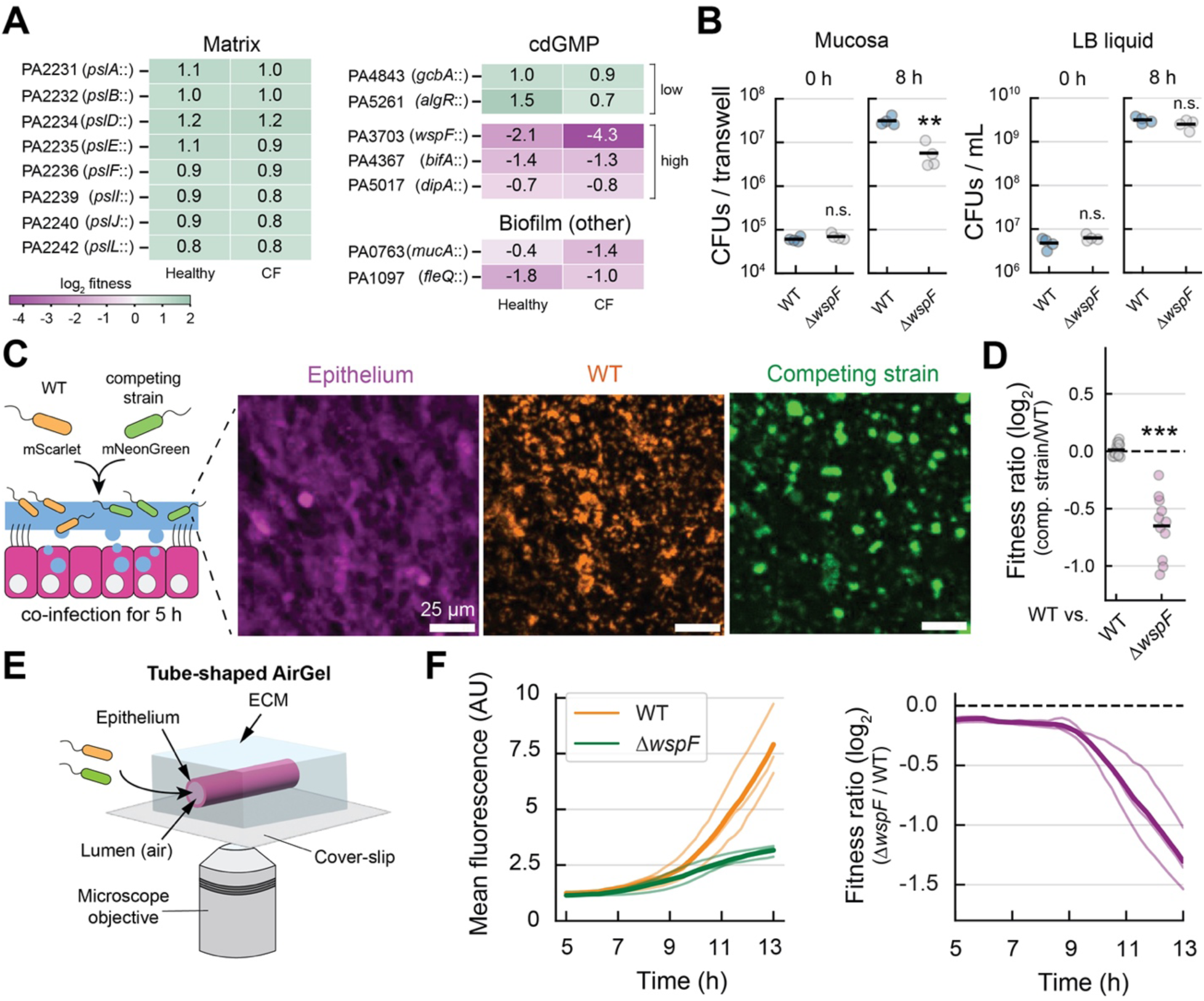
Biofilm formation imposes a fitness penalty during mucosal colonization. **A.** Fitness effects of transposon insertions in representative biofilm-related genes identified by Tn-seq. **B.** CFU-based validation of the fitness changes induced by high cdGMP. Competitions of WT and Δ*wspF* at the mucosa (left) and in LB liquid (right). Each data point represents an independent biological replicate (*n* = 4); the horizontal black lines mark their mean. **C.** Experimental design for imaging of competition in HBE cultures. WT and mutant were mixed 1 to 1, and infections lasted 5 h until microscopic imaging. Representative images of epithelium and bacteria (maximum intensity projection; same field of view). **D.** Fitness ratio based on the quantification of the total area measured from surface coverage by Δ*wspF*-mNeonGreen relative to WT-Scarlet during competition assays at the mucosal surface (HBE cultures). Each data point represents one imaged field of view (*n* = 15 for WT vs WT; *n* = 11 for WT vs. Δ*wspF*) distributed within three distinct biological replicates; the horizontal black lines mark their mean. **E.** Experimental design for imaging of competition assays in HBE grown in AirGels. Strains were mixed 1:1, inoculated in AirGels, and imaged for 13 hours. **F.** Quantification of the mean fluorescence for each strain during AirGel infections (left) and the respective fitness ratios (right). Each thin line represents a biological replicate (i.e., one AirGel, *n* = 3); the thick line is their mean. Statistics: panels B and D, Welch unpaired t-test (** *p* < 0.01, *** *p* < 0.001, n.s. *p* > 0.05).

Biofilm-dwelling cells commit to a sedentary, surface-associated lifestyle. We hypothesized that biofilm formation negatively impacts growth at the mucosal surface by limiting *P. aeruginosa*’s ability to navigate toward profitable environments while locally depleting nutrients. To test this hypothesis, we imaged mucosal surface colonization live at single-bacterium resolution. In competitions between WT and Δ*wspF* constitutively expressing mScarlet and mNeonGreen, respectively, we noticed WT spread uniformly on the mucosal surface while Δ*wspF* remained aggregated into biofilms (Fig. 3C).

Quantification of surface coverage showed that Δ*wspF* could not colonize the mucosal surface to the same extent as WT (Fig. 3D). We also observed a similar spatial distribution when competing WT with Δ*bifA*, another mutant with constitutively high cdGMP levels (Fig. S2) ^43,44^.

To further illuminate the differences in the exploratory behavior of biofilm-dwelling cells, we imaged the dynamics of mucosal colonization in AirGels (Fig. 3E). When competing with WT, Δ*wspF* displayed two successive phases of colonization (Fig. 3F). In an initial phase Δ*wspF* grew exponentially, yet with a lower rate than WT. Later (> 9 h), Δ*wspF* growth stalled, and WT grew dramatically faster. At this stage, Δ*wspF* formed large biofilms that could not efficiently disperse, whereas WT was able to colonize new environments (Movie S1). To understand the mechanisms leading to this growth phase change, we inspected biofilm morphogenetic differences earlier during colonization. WT colonized the mucosal surface into two distinct subpopulations: biofilms and planktonic isolated cells (Fig. 4A). In contrast, Δ*wspF* preferentially formed aggregates while planktonic cells were rare (Fig. 4A). Measuring the surface area of cell clusters in competition experiments shows that Δ*wspF* tends to form larger clusters than WT (Fig. 4B). WT displayed a higher number of small cluster and single cells when compared to Δ*wspF* (Fig. 4C). We conclude that Δ*wspF* cells were mainly forced to stay in biofilms while WT formed looser structure allowing single cells to disperse. We observed similar trends when competing WT with the high cdGMP mutant Δ*bifA* (Fig. S2).

**Figure 4.**
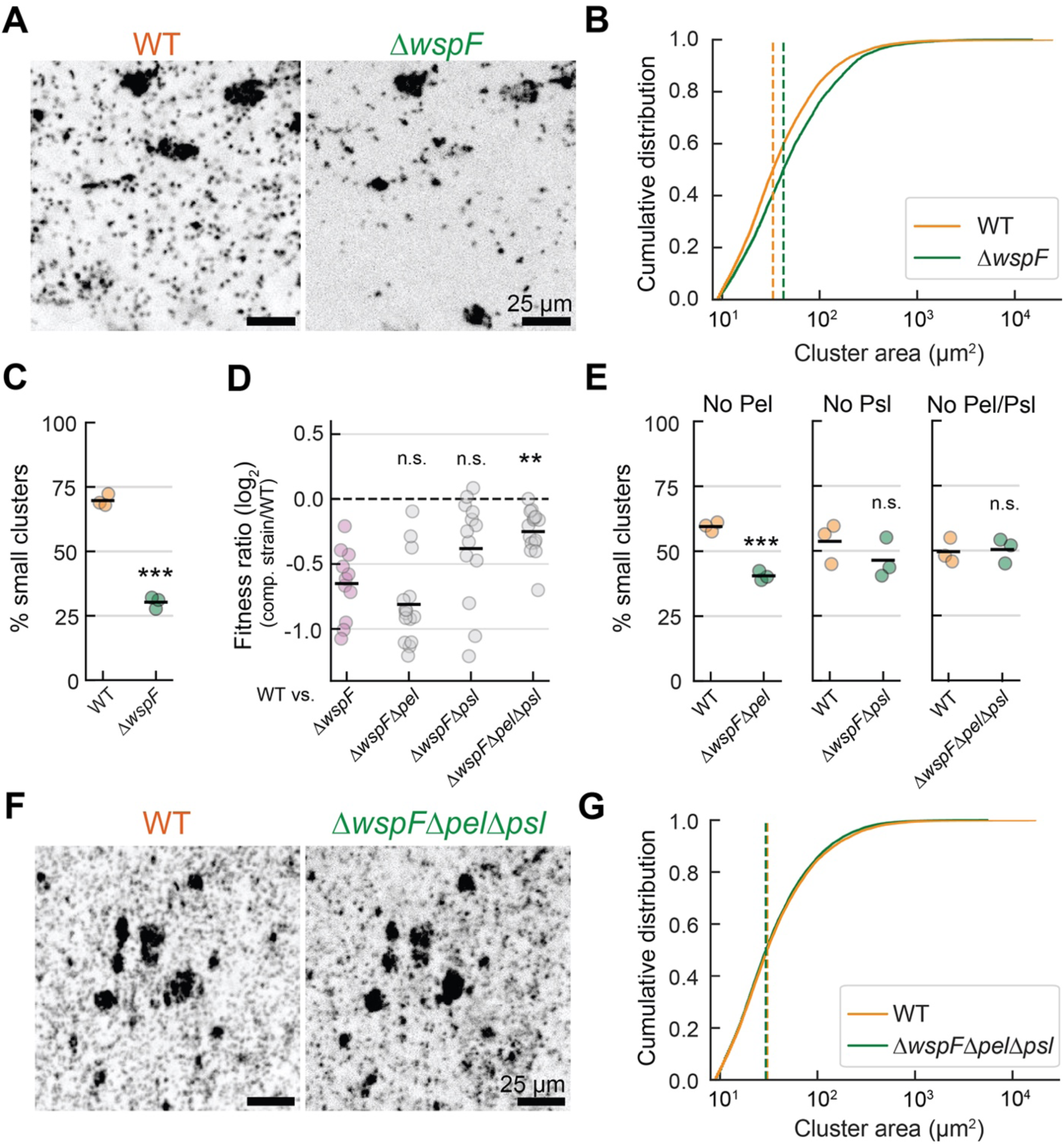
Polysaccharide overproduction causes fitness defects during mucosal colonization. **A.** Representative images of WT and Δ*wspF* competition assays in HBE cultures (maximum intensity projection; same field of view). **B.** Cumulative distributions for cluster sizes formed by WT and Δ*wspF*. The distributions combine three independent replicates. Vertical lines indicate the median cluster-size for each strain. **C.** Percentage of small clusters (smaller than 20 µm^2^) for WT and Δ*wspF* populations. Each data point represents an independent biological replicate (*n* = 3); horizontal black lines mark their mean. **D.** Fitness at the mucosal surface is affected by matrix hypersecretion but not by cdGMP-induced growth defects. Fitness ratio based on relative surface coverage by WT- mScarlet and competing strain expressing mNeonGreen. Each data point represents one field of view (*n* = 11, 15, 13, and 15, respectively) among three biological replicates. Horizontal black lines mark the mean fitness ratio for each condition. **E.** Percentage of small clusters represented by WT and competing strains lacking EPS polysaccharides. Each data point represents an independent biological replicate (*n* = 3); horizontal black lines mark their mean. **F.** Representative images of WT and Δ*wspFΔpelΔpsl* competition assays in HBE cultures (maximum intensity projection; same field of view). **G.** Cumulative distributions for cluster size formed by WT and Δ*wspFΔpelΔpsl*. Distributions include three biological replicates. Vertical lines represent the median cluster-size for each strain. Statistics: panel D, 1-way ANOVA with Tukey HSD multiple comparison test, with asterisks showing significant differences relative to Δ*wspF* competition; panels C and E, Welch unpaired t-test (*** *p* < 0.001, n.s. *p* > 0.05).

In *P. aeruginosa*, matrix components Pel and Psl maintain biofilm cohesion, thereby limiting cell mobility. Δ*wspF and* Δ*bifA* stimulate biofilm production by hyper-secreting Pel and Psl secretion ^13,43^, ultimately reducing mobility, but elevated cdGMP may also cause physiological changes that impact *P. aeruginosa’s* growth at the mucosal surface. To test whether the growth defects were due to a limited ability to explore the epithelial surface, we inspected the fitness of Δ*psl* and Δ*pel* mutants in a Δ*wspF* background. We competed Δ*wspF*Δ*pel*, Δ*wspF*Δ*psl*, and Δ*wspF*Δ*pel*Δ*psl* with WT at the surfaces of HBE cultures.

Δ*wspF*Δ*psl* almost entirely recovered WT-level fitness (Fig. 4D) and produced a number of small clusters similar to WT. Δ*wspF*Δ*pel* had an intermediary phenotype between WT and Δ*wspF* in terms of fitness and amounts of small clusters (Fig. 4D-E). Finally, fitness levels and cluster size distributions of the triple mutant Δ*wspF*Δ*pel*Δ*psl* were indistinguishable from WT (Fig. 4D-G). We conclude that polysaccharide overproduction causes fitness defects, and additional effects from elevated cdGMP levels have a limited impact on the intrinsic fitness of *P. aeruginosa* cells at the mucosal surface.

### Biofilms mechanically damage epithelia while constraining pathogenicity

Planktonic *P. aeruginosa* populations more frequently access epithelia, where they efficiently deploy contact-dependent toxic effectors such as the type III secretion system^46^. We hypothesized that conversely, the sedentary lifestyle of biofilm-dwelling *P. aeruginosa* cells limits their ability to damage tissue. We, therefore, compared HBE cell integrity during long-term AirGel infection by WT and Δ*wspF* (Fig. S3A). WT spreads uniformly over the epithelium surface while rapidly killing epithelial cells, which subsequently undergo lysis (Fig. S3B, Movie S2-3). By contrast, Δ*wspF* biofilms did not lyse epithelial cells to the same extent as WT. Instead, expanding biofilms opened up nodules that stretched out within the tissue (Fig. S3B, Movie S2-3). To understand how these distinct phenotypes impact epithelial viability, we visualized cell death in space and time in AirGels (Fig. S3A). For both strains, host cell death was localized to areas colonized by *P. aeruginosa*. WT had a strong and uniform cytotoxic effect (Fig. S3C, Movie S4). While host cells lysed more slowly during Δ*wspF* infection, the biofilms showed cytotoxic activity towards epithelial cells in the immediate vicinity of the biofilm cell clusters (Fig. S3C, Movie S4). Overall, while biofilm formation negatively impacts *P. aeruginosa* growth by limiting access to nutrients, it may maintain local host tissue integrity over longer timescales. To reach a chronic state, other external pressures must, however, select for biofilm production. The administration of antibiotics during acute infection could exert such pressure on *P. aeruginosa* populations at the mucosal surface.

### Colonization vs tolerance trade-off for biofilms during antimicrobial treatment

To illuminate *P. aeruginosa*’s fitness determinants during antibiotic treatment at the mucosal surface, we again turned to functional genomics in HBE cultures. We performed Tn-seq at the mucosal surface on *P. aeruginosa* populations treated with ciprofloxacin or tobramycin, two clinically relevant antibiotics (Fig. 5A) ^2^. We surveyed tolerance determinants by identifying transposon insertions in genes affecting fitness relative to untreated negative control (Fig. S4A). Mutations in 45 genes affected tolerance to ciprofloxacin and 235 genes to tobramycin (Table S4). We identified that mutations in *nfxB* and *mexZ* increase fitness to ciprofloxacin and tobramycin, respectively (Fig. S4B). This validated our screen as these mutants overexpress the efflux systems MexCD-OprJ and MexYX, which induce resistance to ciprofloxacin and tobramycin, respectively ^47–49^. We found a mild negative correlation between fitness mucosal growth fitness and ciprofloxacin treatment (Fig. 5B). This is consistent with the mode of action of ciprofloxacin that requires growth to kill bacteria ^50^. Mutants displaying fitness defects during mucosal colonization generally showed increased tolerance to ciprofloxacin (Fig. S4B). There was, however, no correlation between mucosal and tobramycin-induced selections (Fig. 5B), which may be explained by tobramycin’s more intricate mode of action that does not imperatively rely on growth ^50,51^.

**Figure 5.**
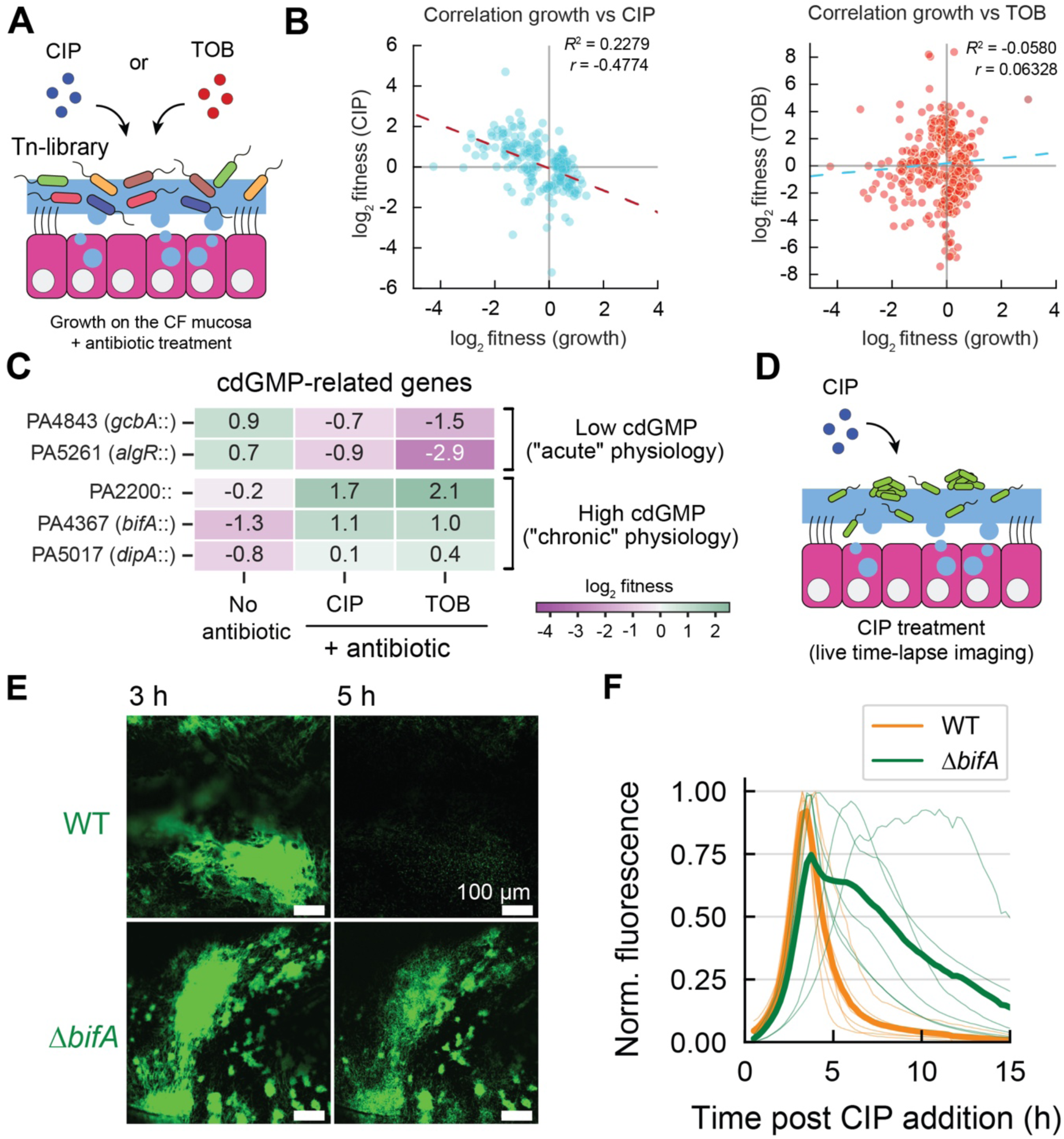
Colonization vs tolerance trade-offs for biofilms during antimicrobial treatment. **A.** Schematic representation of the Tn-seq experiment in the presence of antibiotics at the mucosal layer. **B.** Correlation of fitness of *P. aeruginosa* transposon mutants in the presence and absence of antibiotics in HBE cultures. Each data point represents a gene found in the Tn-seq. Dashed lines show a linear fit of the data. *r*: Pearson correlation coefficient. *R*^2^: coefficient of determination. **C.** Fitness effects of transposon insertions in representative cdGMP-related genes. Genes shown have passed the significance cutoff (p < 0.05) in at least one condition. **D.** Simplified representation of time-lapse imaging in AirGels upon CIP treatment. **E.** Representative images of WT and Δ*bifA* after 3 h and 5 h of CIP exposure in AirGels (average intensity projection). **F.** Fluorescence-based quantification of WT and Δ*bifA* tolerance to CIP in AirGels (min-max normalized). Thin lines represent one independent biological replicate (*n* = 6), and thicker lines are their normalized mean.

In addition to compound-specific selection, 24 mutants were tolerant to both antibiotics (Table S4). Among them, mutants involved in cdGMP signaling genes were under strong selection (Fig. 5C, Table S4). Opposite to mucosal colonization, mutations increasing cdGMP levels improved fitness, while ones reducing cdGMP decreased fitness (Fig. 5C). This is consistent with the expectation that the biofilm lifestyle metabolically and physically reduces the efficacy of antimicrobials ^15,16^. To better expose the biophysical mechanisms of tolerance, we used live imaging to compare the sensitivities of biofilms and planktonic cells to ciprofloxacin. We infected AirGels with WT and Δ*bifA*, allowed the strains to colonize the mucosal surface, and subsequently introduced ciprofloxacin. We compared their tolerance to treatment based on their respective fluorescence intensity changes (Fig. 5D). Overall, Δ*bifA* biomass decreased more slowly than WT (Fig. 5E-F, Movie S5), confirming its increased tolerance and demonstrating that biofilms protect *P. aeruginosa* from antimicrobials during AirGel infections. Thus, we identified a trade-off for biofilm formation at the mucosal surface: forming biofilms penalizes *P. aeruginosa* but confers protection against antibiotic treatment. This model provides a rationale for *P. aeruginosa* evolution *in vivo,* which manifests itself in transitions from acute to chronic states. Despite these clear evolutionary pressures, clinical isolates from chronically infected patients do not always show a clear antibiotic-tolerant, hyper-biofilm phenotype, even under prolonged and aggressive antibiotic administration ^10,52^. This motivated us to next explore the benefits of strain diversity and genotypic diversity for fitness.

### Biofilms cross-protect planktonic cells from antibiotics

In human patients, diversification leads to different clones co-existing within the same lung region, including acute/planktonic and chronic/biofilm mutants ^10,53,54^. We thus wondered whether population heterogeneity improves fitness to otherwise antibiotic-sensitive clones. We hypothesized that chronic strains protect acute planktonic ones within biofilms. To test this hypothesis, we co-infected AirGels with WT (acute-like) and Δ*wspF* (chronic-like), each expressing a distinct fluorescent protein. We imaged these populations while treating them with ciprofloxacin (Fig. 6A). Before treatment, *P. aeruginosa* Δ*wspF* formed the largest biofilms, while WT cells occasionally formed smaller clusters. Oftentimes, WT co-colonized large Δ*wspF* biofilms (Fig. 6B)

**Figure 6.**
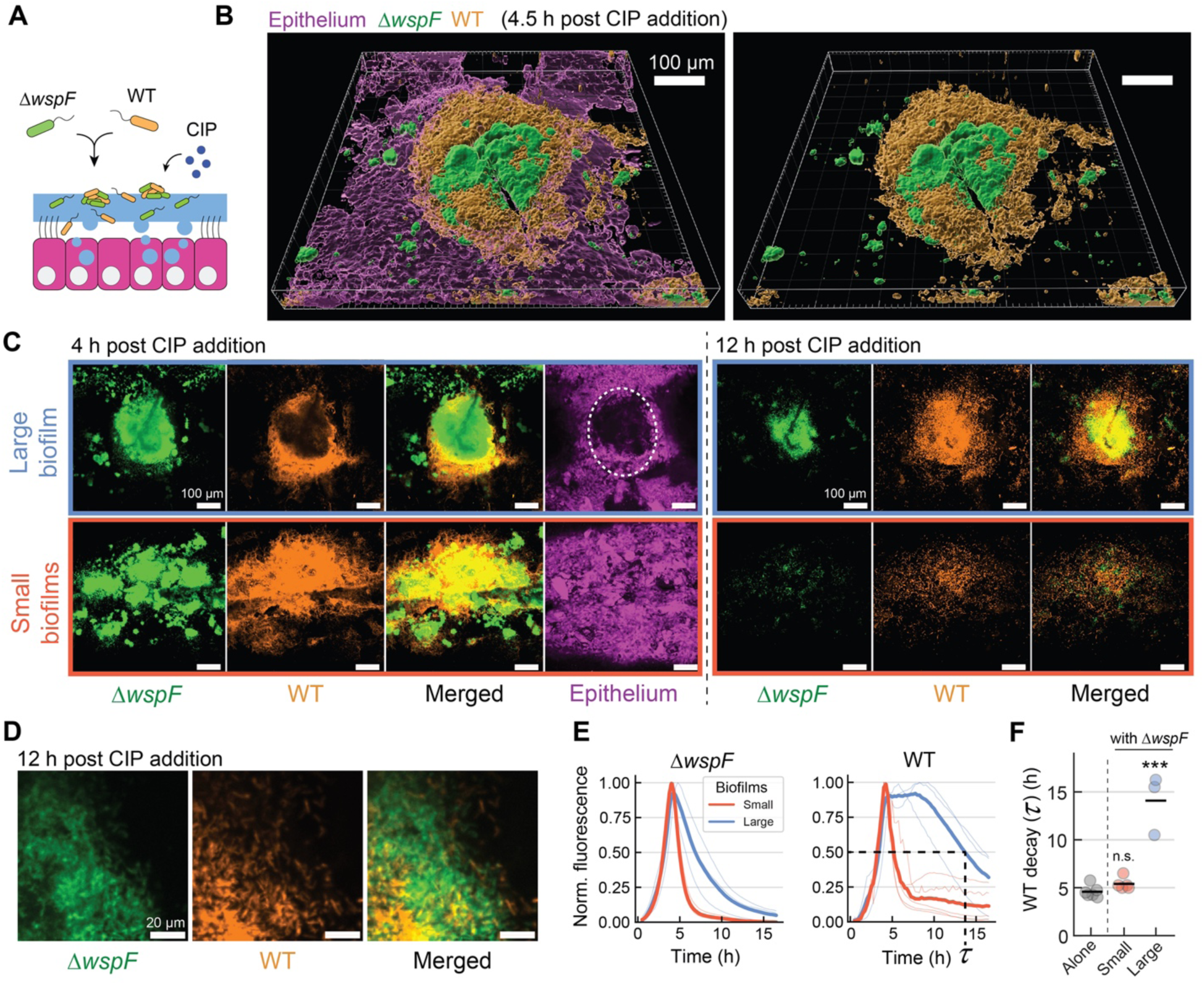
Biofilms cross-protect planktonic cells from antibiotics. **A.** Simplified representation of time-lapse imaging of competitions in AirGels upon CIP treatment. **B.** 3D rendering of a large Δ*wspF*-WT mixed *P. aeruginosa* biofilm. The right image only shows *P. aeruginosa* strains without epithelium. **C.** Representative images illustrating differences in tolerance of mixed infections where Δ*wspF* forms large or small biofilms (maximum intensity projection). WT often co-colonizes large Δ*wspF* biofilms. After 4 h of treatment, Δ*wspF* created nodules stretching out within the tissue (white circle). The two distinct regions shown were collected in the same AirGel. **D.** Representative confocal images showing WT cells surviving within Δ*wspF* biofilms after 12 h of CIP treatment. **E.** Fluorescence-based quantification of tolerance to CIP of regions containing small or large biofilms (min-max normalized). The categorization of “large” or “small” is based on the presence of Δ*wspF* biofilms large enough to form nodules within the tissue. Each thin line represents one field of view (*n* = 3 for large biofilms; *n* = 5 for small biofilms) distributed among two AirGels. Thick lines mark the mean normalized fluorescence. We computed the time at which the WT normalized signal decreased to 50% from its maximum value (*τ*, or “WT decay”) within each replicate. **F.** WT decay *τ* upon CIP treatment for WT alone, WT in the presence of small Δ*wspF* clusters, or WT in the presence of large *ΔwspF* clusters. Statistics: panel F, 1-way ANOVA with Tukey HSD multiple comparison test, with asterisks showing significant differences relative to “WT alone” condition (*** *p* < 0.001, n.s. *p* > 0.05).

We next quantified the population dynamics upon ciprofloxacin exposure. WT planktonic cells rapidly died upon treatment, as were small clusters of WT and Δ*wspF* (Fig. 6C). By contrast, large Δ*wspF* biofilms were more tolerant (Fig. 6C). In addition, WT cells growing in or near large Δ*wspF* biofilms were killed more slowly than the rest of the WT population (Fig. 6C-D). To reveal this cross-protection mechanism, we compared WT survival in regions containing large versus small Δ*wspF* cell clusters. WT cells survived much longer within large Δ*wspF* biofilms than in regions that only contained small clusters (Fig. 6E-F, Movie S6-7). Biofilms, therefore, shield planktonic cells from antibiotics, potentially allowing them to grow again when treatment stops and potentially exacerbate cytotoxicity. Biofilm-dependent cross-protection of acute strains may thus contribute to unexpected evolutionary paths during chronic infections. However, such an effect is only relevant when chronic *P. aeruginosa* has time to form sufficiently large biofilms, highlighting the importance of early therapeutic interventions.

## Discussion

*P. aeruginosa* takes on complex evolutionary paths when transitioning from acute to chronic phases of infections ^19,55,56^. Here, we combined functional genomics with a tissue-engineered lung model to decode how *P. aeruginosa* physiologically adapts to the airway mucosal environment. Leveraging human organoids allowed us to better emulate the chemical and physical conditions the pathogen experienced during colonization. Investigating *P. aeruginosa* physiology in a mucosal-like model revealed the conflicting biophysical factors that impact its fitness during infection. These tradeoffs likely favor phenotypic heterogeneity and flexibility, which could allow *P. aeruginosa* to persist during different stages of infection.

At the metabolic level, we found that *P. aeruginosa* heavily relies on sugars and lactate, both of which are available in the lungs of infected patients ^27,28,57^. In addition, *P. aeruginosa* compensates for the scarcity of amino acids and nucleotides at the surface of HBE cultures. *P. aeruginosa*’s metabolic adaptation is thereby reminiscent of the strategies adopted by other microbes to the mucin-rich environments of their hosts ^58,59^. Metabolic independence improved fitness at the early steps of infection, but the metabolic pressure could be relaxed at later stages. For example, sputum often contains copious amounts of dead host cells, which can constitute a pool of amino acids ^27,28,30,60–62^. Those cells, often neutrophils killed by *P. aeruginosa*, are not recapitulated in our infection model. In fact, our models lack long-term inflammation features fueled by chronic infection ^63,64^. Therefore, while we found that mucins are an important nutrient source and *de novo* amino acid and nucleotide biosynthesis are required early on, incorporating immune cells could impact *P. aeruginosa’s* metabolic state on longer timescales.

Our results provide a new perspective on *P. aeruginosa*’s *in vivo* evolution towards chronicity. Genomic studies based on human clinical isolates have identified the mutational paths leading to *P. aeruginosa*’s chronic lifestyle. In these conditions, mutations inducing loss of motility and acute virulence factors (such as T3SS), along with increased biofilm formation and antibiotic resistance, appear to be selected for ^7,34^. Thus, our approach contributes to our understanding of the forces and mechanisms driving *in vivo* evolution. In particular, we found that biofilm genes, including those linked to cdGMP signaling, were critical determinants of colonization. Biofilm formation controlled via cdGMP signaling incurs a strong fitness trade-off between mucosal colonization and antibiotic tolerance during infections (Fig. S5). While inhibiting effective mucosal colonization, strong biofilm formation protects cells against antibiotic treatment. Consistent with this, in isolates of chronically infected patients, *bifA* and *wsp* operon genes have been identified as evolution hot spots ^19,65^. Therefore, our work highlights evolutionary pressure on the genes controlling cdGMP levels in the clinical setting.

In chronic lung infections, although dominated by *P. aeruginosa* strains with hyper-biofilm phenotypes, planktonic strains with high cytotoxicity often re-emerge ^53–55^. These rebounds have been suggested to play a crucial role during the development of pulmonary exacerbations leading to a dramatic decline in lung function ^66–68^. We showed that a chronic strain can accommodate cells from acute strains within their biofilm. As a result, chronic strains shelter sensitive mutants from antibiotic treatment. This protection mechanism could thereby prevent the eradication of acute strains during treatment, constituting a potential reservoir for re-colonization after the antibiotic is removed (Fig. S5). Phenotypic diversification within biofilms may represent a general protection strategy against strong selective pressures. For example, cells in biofilms avoid predation from immune cells or from bacteriophages ^7,69–73^. Cross-protection, therefore, factors in the design of phage therapies for chronically infected CF patients ^74,75^.

Antimicrobial treatment failure is one of the most pressing public health challenges ^17,50^. Developing technologies that probe pathogens under physiologically-relevant conditions will allow us to directly identify their vulnerabilities. We showed that combining organoid models with omics-based methods can resolve essential molecular mechanisms of infections ^31^. Human organoids as infection models are potentially widely applicable to other pathogens as well as more complex microbial communities, including microbiota. The modular design of tissue-engineered systems will enable expansion to other sensitive organs and the incorporation of additional selective pressures such as immune components ^76,77^. This approach could be instrumental in the development of innovative treatments against resistant pathogens that target alternative aspects of their physiology.

## Materials and methods

### General bacterial culturing

Media for bacterial culturing used Luria-Bertani (LB) broth and agar (1.5%), supplemented with antibiotics whenever necessary. Antibiotics were prepared in concentrated stock solutions (100x or greater) and stored at -20°C until use. Ciprofloxacin was dissolved in 20 mM HCl, while gentamicin, tobramycin and tetracycline were dissolved in sterile deionized water. Unless mentioned otherwise, all liquid cultures were incubated at 37°C with shaking at 225 rpm.

### Bacterial strain construction

We used *P. aeruginosa* strain PAO1 for all our experiments ^78^. A complete list of strains, plasmids, and primers used can be found in Table S5.

To generate clean in-frame deletions, approximately 750 to 1000-bp fragments immediately upstream and downstream of the target locus were cloned using Gibson assembly into the pEX18Gm suicide vector ^79,80^. Fragments amplified from *P. aeruginosa* PAO1 genomic DNA (gDNA) and cleaned up using the Monarch PCR & DNA Cleanup kit (New England Biolabs) were used for Gibson assembly together with pEX18Gm cut with EcoRI-HF and Xbal. The assembled construct was then transformed into *E. coli* XL10- Gold (Agilent), with transformants selected on LB agar plates with 10 µg/mL gentamicin. Correctly assembled plasmids were identified by colony PCR and verified by Sanger sequencing (Microsynth, Switzerland). The constructs were then miniprepped (using GeneJET Plasmid Miniprep kit, Thermo Scientific) and transferred to *E. coli* S17 ^81^ by electroporation, the strain used as a donor during biparental conjugations. Biparental conjugations were performed following previously described protocols ^82^, leading to the insertion of the constructs into the *P. aeruginosa* PAO1 genome. Merodiploids were selected on a VBMM medium with 60 µg/mL gentamicin ^82^. Then, to select colonies resulting from homologous recombination, the merodiploids were plated on LB lacking NaCl and containing 10% sucrose. Colonies containing the unmarked deletions were identified by PCR and verified by Sanger sequencing. All the primers used, and the resulting strains can be found in Table S5.

Finally, fluorescent strains used for microscopy experiments were generated using previously published plasmids containing the fluorescent proteins mScarlet or mNeonGreen under the control of the *tet* promoter, which is constitutive in *P. aeruginosa*. The respective construct was inserted in the genome of PAO1 at the *att*Tn7 by electroporation, followed by selection in LB with 60 μg/mL gentamicin ^26,82^.

### Transposon library preparation

We conjugated the Tn*5*-based transposon T8 (IS*lacZhah*-tc) into *P. aeruginosa* PAO1 as previously described ^83,84^. In summary, *P. aeruginosa* PAO1 WT was grown overnight in liquid (five cultures containing 5 mL each) at 42 °C without shaking. At the same time, five plates of *E. coli* SM10λ*pir* (carrying the plasmid pIT2 containing the transposon) were grown overnight in LB-agar containing 100 μg/mL of carbenicillin. These five plates of *E. coli* SM10λ*pir* were harvested (5 mL total), washed in LB, and resuspended to an OD600 of ∼160. Next, the overnight cultures of *P. aeruginosa* PAO1 WT (recipient) were washed and resuspended in aliquots of 50 µL containing an equivalent of an OD600 of 50. Donor and recipient were mixed (roughly 3:1 ratio in OD600), spotted into 0.22 µm filter discs pre-incubated on top of LB plates (50 µL per filter, total of 20 filters), and incubated 2.5 h at 37°C. The 20 conjugation spots were pooled and resuspended into 16 mL of LB. Finally, to select for transposon PAO1 transposon insertion mutants, 70 aliquots (200 µL each) were plated on large (150 by 25 mm) LB agar plates containing tetracycline (60 µg/ml) and chloramphenicol (10 µg/ml). The plates were incubated for 24 h at 37°C plus 18 h at room temperature. After incubation, colonies from all the plates (∼3,500 per plate) were resuspended in LB containing 15% glycerol and stored as 500 µL aliquots at -80 °C. The final library stock had an OD600 of ∼56, consisting of ∼250,000 unique mutants. These estimations are based on the number of pooled colonies and the number of unique insertions identified by sequencing.

### Tn-seq experimental design (mucosal growth and antibiotic tolerance)

We performed two distinct Tn-seq experiments: (i) a screen searching for *P. aeruginosa* genes important for growth on the mucosal surface (Fig. 1) and (ii) a screen looking for genes involved in antibiotic tolerance under these conditions (Fig. 5).

<u>Tn-seq for growth at the mucosal surface:</u> the library stock was diluted in LB to an OD600 of 0.25 (3 mL) and grown until deep stationary phase for 14 h. Then, cells were spun down, washed in PBS buffer, resuspended in PBS at an OD600 of 1, and used as inoculum for HBE infections. Mucosal epithelia developed from HBE cells derived from a non-CF and a CF donor were used during these infections. Ten different Transwell inserts for each donor were infected using 1 µL, which represents ∼10^6^ *P. aeruginosa* cells per insert. The infection progressed for 11 h, which resulted in ∼3 generations without invasion of the tissue by the library. Importantly, each Transwell insert contained HBE cells that differentiated for around 2 months without any washes above the epithelium, which ensured that abundant amounts of mucus were produced. We used stationary phase cells for infections to force *P. aeruginosa* to adapt and grow exclusively from nutrients found in the mucosal environment. After infection, the tissue was homogenized using 100 µL of Triton X-100 (0.1% in PBS). Each insert was vigorously scraped until all the tissue material was removed. Then, samples from the ten distinct inserts were pooled together, vortexed and pipetted vigorously, spun down (2 min, 14,000 rpm), washed in PBS to remove traces of Triton, spun down again, and the pellets were stored at -80 °C until subsequent processing. To determine the exact inoculum size and number of generations during infection, the inoculum and an aliquot of the stored pellet were platted for CFUs.

In parallel, we prepared an LB control sample in which the inoculum used during the infections was grown in LB for the same amount of time as the infections (Fig. S1B). For these, the inoculum was diluted in LB (3 mL) for an OD600 of 0.1. The culture was grown in parallel with the infections for 11 h. Then, 1 mL of the culture was spun down, resuspended in Triton X-100 (0.1% in PBS) and incubated for 15 min, spun down, washed in PBS, spun down again, and the pellet was stored at -80 °C until subsequent processing. To determine the exact inoculum size and number of generations during growth, we platted the culture for CFUs at the start and the end of the experiment.

<u>Tn-seq for tolerance at the mucosal surface:</u> the infection was set up identically to the infection Tn-seq above, but 30 Transwell inserts with differentiated epithelia from CF HBE cells were infected with the inoculum. After 11 h of infection, these inserts were split into 3 conditions: (i) ciprofloxacin treated, (ii) tobramycin treated, and (iii) untreated negative control. Each treatment used 10 inserts. For ciprofloxacin treatment, the antibiotic was added to the bottom and top of the insert at a concentration of 1 µg/mL, and exposure lasted 20 min. For tobramycin treatment, a concentration of 50 µg/mL with exposure lasting 30 min. After antibiotic treatment, tissue was homogenized using 100 µL of Triton X-100 (0.1% in PBS) as described before. Then, samples from the 10 distinct inserts within each treatment were pooled together, vortexed and pipetted vigorously, spun down (2 min, 14,000 rpm), and washed in PBS to remove traces of Triton and the antibiotics. The same protocol was applied for the untreated negative control. Each of the pellets was then resuspended in 3 mL of LB, followed by an outgrowth step of 7 to 12 hours, depending on the treatment, leading to an OD600 of 1.7-2.2. 1 mL of each culture was spun down, and the pellet was stored at -80 °C for later processing. CFUs were plated at every step for monitoring of survival and growth. Ciprofloxacin and tobramycin treatments lead to a survival rate of ∼1%.

### Library preparation and sequencing

gDNA was extracted from the samples using the QIAamp DNA Mini kit (Qiagen). Library preparation and sequencing were performed at the Lausanne Genomic Technologies Facility, located at the University of Lausanne. In summary, gDNA (500 ng) was first sheared with a Covaris S220 using 400 bp insert settings (50 µL in microTUBES with AFA fiber, Peak incident power: 175, Duty factor: 5%, Cycles per burst: 200, time: 70 sec). After a purification with SPRI beads at a 1.6x ratio, libraries were prepared with the xGen DNA MC UNI Library Prep Kit (IDT, protocol version v2) using xGen UDI-UMI adapters (IDT, 15 µM stock). With these adapters, P5 and P7 sequences are inverted compared to Illumina adapters, allowing transposon sequencing directly from read 1 (P5 side) in a single-end run. The purified ligated product was PCR amplified with a primer specific for the Illumina P7 sequence (CAAGCAGAAGACGGCATACGA) and a second one specific for the transposon sequence (cgacgttgtaaaacgaccacgt) carrying a 5’-biotin. PCR was performed with the KAPA HiFi HotStart ReadyMix kit (Roche). Cycling conditions were 98 °C for 45 s, followed by ten cycles of 98 °C for 15 s, 60 °C for 30 s, and 72°C for 30 s, and a final extension of 1 min at 72°C. The library was purified with SPRI beads at a 1x ratio.

The PCR product was captured with pre-washed Dynabeads MyOne Streptavidin T1 (Thermo Fischer). Binding & Wash (B&W) Buffer 2x composition is 10 mM Tris-HCl pH 7.5, 1 mM EDTA, 2M NaCl. At least 1 ml of 1x B&W Buffer was added and mixed with 25 µl of Dynabeads. After 1 min on a magnet, the supernatant was removed, and Dynabeads were washed with the same volume of 1x B&W Buffer. This wash was repeated a second time. After removal of the supernatant on the magnet, the Dynabeads were resuspended with 50 µl of 2x B&W Buffer. 50 µL of the library was mixed with the washed Dynabeads, followed by incubation at RT on a rotator for 30 min. After 2 min on a magnet and discarding the supernatant, a wash was done with 100 µL of 1x B&W Buffer. Two additional washes were done before the final elution in 40 µL H2O.

Half of the washed capture was used for the nested PCR with the Illumina P7 sequence (see sequence above) and a tailed primer made of the Illumina P5 sequence (red), the TruSeq read 1 primer binding site (black), and a transposon specific binding sequence (orange) (Table S5). This nested PCR amplification was performed with the KAPA HiFi HotStart ReadyMix kit with the same cycling conditions as above but eight cycles. The final library was purified with SPRI beads at a 0.9x ratio. It was quantified with a fluorometric method (QubIT, Thermo Scientific), and its size pattern was analyzed with a fragment analyzer (Agilent).

Sequencing was then performed on a NovaSeq 6000 (Illumina) on a lane of an SP flow cell for a 100-cycle single-end sequencing run. Clustering was performed with a 1.2 nM library spiked with PhiX (Illumina). Sequencing data were demultiplexed using the bcl2fastq conversion software (version 2.20, Illumina) and further processed for transposon insertion analysis (see next).

### Tn-seq data processing and analyses

For each sample, demultiplexed raw reads originating from two distinct lanes were first merged into a single sequencing file. Then, sequencing processing and mapping to the *Pseudomonas aeruginosa* PAO1 genome (NCBI: NC_002516) was done using TPP (with BWA for mapping) ^85,86^. Parameters included: “Tn5” as the protocol, “mem” as the algorithm, and other default options provided by software (e.g., “Max reads” = -1, “Mismatches allowed” = 1, “Start window to look for prefix” = 0,20). Next, we used TRANSIT v3.2.7 ^85^ to perform quality control and further processing with the “wig” files generated by TPP. Because our dataset showed high skewness, we normalized all the samples using the Beta-Geometric Correction (BGC) as recommended by the software’s manual (command “normalize” with -n aBGC). Finally, we used the Tn5 “resampling” method within TRANSIT (GUI mode) to assess the conditional essentiality of genes between two conditions in five distinct comparisons (Fig. S1B; Fig. S4A). Within each comparison, the parameters followed default conditions (samples=10000, pseudocounts=0.00, adaptive=True, histogram=False, includeZeros=True), except no normalization was used since wig files had already been normalized. Specifically, the following comparisons were tested, always with the first sample used as the “control sample” and the second as the “experimental sample”: (1) inoculum vs. healthy HBE; (2) LB control vs. healthy HBE; (3) LB control vs. CF HBE; (4) no antibiotic vs. ciprofloxacin treated; (5) no antibiotic vs. tobramycin treated. Comparisons 1-3 are related to mucosal growth Tn-seq, and comparisons 4-5 are related to the tolerance Tn-seq. The results of these comparisons are available in Tables S1, S2, and S4. Notably, even though we did not have individual sequencing replicates, each sample in our dataset consisted of a pool of material from ten different Transwell inserts (see the Tn-seq experimental design session above), which already accounts for biological variability. Finally, as suggested by the software, we used the “adjusted *p*-value” < 0.05 as the cutoff for statistical significance within each comparison (which applies the Benjamini-Hochberg method for controlling for the false discovery rate).

As additional controls, we also ran the resampling method with normalization by the default Trimmed Total Reads (TTR). Overall, genes that passed the significance cutoff for the BGC normalized data also did so for the TTR. However, TTR normalization led to a much larger number of genes passing the significance cutoff, including several cases where the log2 fold-change difference was small. Because it has been shown that BGC normalization reduces false positives derived from Tn-seq data skewness ^87^, we used the more conservative results derived from the BGC normalization. Tn-seq sequencing data will be deposited at GEO upon acceptance of the manuscript.

### Additional Tn-seq-related data processing and analyses

#### Functional classification using PseudoCAP

For the classification of gene functions shown in Fig. 5C, we used the Pseudomonas Community Annotation Project (*PseudoCAP*) functional classification ^88^. We counted the number of times each term appeared in the list of 620 genes identified as significant (p < 0.05) while *P. aeruginosa* grew on HBE cultures produced from the primary cells of a non-CF donor. The control sample here was the culture used as inoculum for the infections (Fig. S1B). We did not include genes classified as hypotheticals or coding for non-determined “putative enzymes.” Also, to avoid redundancy, we renamed the classification for genes following in both “Membrane proteins” and “Transport of small molecules” categories to “Membrane transport”.

#### Correlation analyses

Analysis in Fig. 1D involved genes identified while *P. aeruginosa* grew on HBE cultures produced from the primary cells of a non-CF and a CF donor. The control sample here was the LB culture grown for the same time as the infection progressed (Fig. S1B), which allowed us to narrow down the list of genes identified and filter out genes likely related to broad growth defects. Genes shown as significant (*p* < 0.05) by the analysis with TRANSIT in at least one of the two samples were used to make the plot. Analyses shown in Fig. 5B involved Tn-seq data for antibiotics tolerance (ciprofloxacin or tobramycin) compared to the Tn-seq results from growth on CF HBE. This is because the antibiotic tolerance Tn-seq was performed in the same CF cells (see above). Linear regressions and the calculation of the shown Pearson correlation coefficient (*r*) and the coefficient of determination (*R*^2^) were performed using Pandas ^89^ and scikit-learn ^90^ in Python ^91^.

### General tissue culturing for Transwell HBE cultures and AirGels

Human Bronchial Epithelial cells were obtained from Lonza (21TL200815 for healthy donor, 00196979 for the CF donor) and cultured in T-25 flasks using PneumaCult–Ex Plus medium (P-Ex Plus) (Stemcell Technologies). Unless mentioned otherwise, incubations were always done at 37 °C with 5% CO2 using a humidified cell culture incubator. We always limited the expansion of cells to a maximum of three passages. Upon reaching confluence, the cells were detached from the flask using the Animal Component-Free Cell Dissociation Kit (Stemcell Technologies), centrifuged, resuspended in P-Ex Plus, and loaded into Transwells or AirGels, as previously described ^26^ with the minor adaptations highlighted below.

For culturing in Transwells, HBE cells were loaded on tissue culturing inserts with 0.4- μm pore polyethylene therephthalate (PET) transparent membranes (Sarstedt). Loading used 30,000-50,000 cells per well, followed by expansion until confluence with P-Ex Plus (1-3 days) and transition to ALI culture conditions, where the PneumaCult-ALI medium (P-ALI) (Stemcell Technologies) was used on the basal side and air on the apical side. Differentiation proceeded for at least 30 days, with inserts often maintained longer (as specified in the experiment description) for mucus accumulation. Experiments using Transwells involved the Tn-seq (Fig. 1; Fig. 2A; Fig. 2C-D; Fig. 3A; and Fig. 5A-C), the validations with CFUs (Fig. 2B and Fig. 3B), and the competition assays and cluster size quantification (Fig. 3C-D and Fig. 4).

AirGels chip fabrication followed our previous description ^26^. Briefly, a 3D-printed mold was used for cast PDMS chips that served as the AirGel scaffold and PDMS rods to pattern the lumens ^26,92^. AirGel chips were always mounted on glass-bottom 6-well black plates suitable for microscopy (1.5 coverslip) (IBL). The chips were filled with an extracellular matrix hydrogel containing a 1:3 ratio of Matrigel (Corning) and high-density rat tail type I collagen (Corning), followed by a polymerization step at 37 °C with 5% CO2. Rods were then removed, giving the tube shape of AirGels, followed by a crosslinking step. Then, HBE cells were loaded into AirGels using 10-12 µL of cell suspension containing ∼20,000 cells/µL, as done before ^26^. All of our AirGels experiments were done with HBE cells derived from the CF donor (see above). For full protocol on AirGel fabrication, hydrogel preparation, and cell loading, see Rossy et al. ^26^. Upon loading, HBE cells were expanded in AirGels P-Ex Plus until reaching confluence (typically ∼3 days). After confluence was reached, the expansion medium was replaced with PneumaCult Airway Organoid Differentiation Medium (P-organoid) (Stemcell Technologies) supplemented with 5 µM of the protease inhibitor GM6001 (InSolution GM6001, Merck). AirGels were differentiated under immersed conditions (i.e., with media filling both basal and apical sides) for at least one month, with the medium being replaced every Monday, Wednesday, and Friday. Finally, AirGels were transitioned to ALI conditions (i.e., lumen filled with air) 2-7 days before being used on infection experiments. Experiments using AirGels involved the competition assays with dynamic temporal imaging (Fig. 3E-F), the time-lapses assessing *P. aeruginosa* cytotoxicity (Fig. S3), the single-strain antibiotic tolerance assays (Fig. 5D-F), and multi-strain antibiotic tolerance assays (Fig. 6).

### Validations experiments with CFUs

We performed two different experiments with colony-forming units (CFUs) to validate our Tn-seq during mucosal growth: (i) genes involved in metabolism (Fig. 2B) and (ii) the strongest hit related to biofilm (Fig. 3B).

For the validation of genes involved in metabolism shown in Fig. 2B, we picked *edd*, *lldP,* and *pvdE* as these displayed a range of fitness phenotypes and are distinct from genes previously identified in Tn-seq screen of *P. aeruginosa* growing on mucins ^58^. The tested strains (WT Δ*edd*, *lldP,* and *pvdE*) were grown in LB broth from an LB plate for 6 h until reaching dense cultures; then, cells were diluted to an OD600 of 0.25 and grown to a deep stationary phase for 11 h. Next, cells were spun down, washed in PBS buffer, and resuspended in PBS at an OD600 of 0.3, which was used as inoculum for infections. For the determination of the exact inoculum size, each inoculum was plated for CFUs before the infection started. Then, for each tested strain, three distinct Transwell inserts were infected using 1 µL of inoculum, and the infection progressed for eight hours. Each Transwell insert contained tissues from CF HBE cells that had been under differentiation for around three months without any washes above the epithelium, which ensured a large amount of produced mucus (as done during the Tn-seq). After infection time ended, the tissue was homogenized using Triton X-100 (0.1% in PBS) and plated for CFUs. In parallel to the infections, the growth of each strain in LB broth control was also determined. From the same inoculum, three LB cultures were started for each strain at an OD600 of 0.03 (600 µL each). Cultures were grown for the same eight hours as the infections, with CFUs platting at the beginning and end of the experiment. Next, we determined the growth rate (shown in Fig. 2B) based on the number of doublings detected for each condition during the eight hours of the experiment (# generations/8 hours).

For the validation of genes involved in biofilm formation shown in Fig. 3B, we picked *wspF* as it displayed one of the strongest effects in our dataset. In this case, we competed WT and Δ*wspF*, which is possible due to the differences in the colony morphology of these strains (smooth for WT and rugose for Δ*wspF*) ^13^. WT and Δ*wspF* strains were grown overnight in LB broth from an LB plate; then, cells were diluted 1:100 (WT) or 1:50 (Δ*wspF*) in new LB cultures and grown for around 3 h. Next, cells were spun down, washed in PBS buffer, resuspended in PBS at an OD600 of 0.3, and mixed into a 1:1 ratio that was used as inoculum for infections. The inoculum was plated for CFUs before the infection started. Then, four distinct Transwell inserts were infected using 1 µL of the 1:1 inoculum mix, and the infection progressed for eight hours. Each Transwell insert contained tissue from CF HBE cells under differentiation for around 2.5 months. The excess mucus was washed before the infection by overnight incubation with P-ALI media on the top of the Transwell, followed by complete aspiration 4 h before the experiment started. After infection time ended, the tissue was homogenized using Triton X-100 (0.1% in PBS) and plated for CFUs. In parallel to the infections, the growth of the inoculum mix in LB broth was also determined. From the same inoculum, four LB cultures were started for each strain at an OD600 of 0.03 (600 µL each). Cultures were grown for the same 8 h as the infections, with CFUs platting at the beginning and end of the experiment. We counted the amount of WT and Δ*wspF* CFUs in each of these mixed competition assays based on their morphological differences.

### Metabolic modeling, in silico gene essentiality analysis, and dimensionality reduction

To simulate the metabolism of *P. aeruginosa*, we used a published genome-scale metabolic model (GEM) PAO1 ^39^, for which we estimated thermodynamic properties ^93–95^ for 60% of the reactions and 68% of the metabolites. The GEM includes 1,149 genes, 1,147 which are part of the Tn-seq mutant screen. We used the thermodynamically curated to assess the effect of the individual deletion of each one of the genes in the GEM for 61 different media compositions. For the analysis we used complex media broadly utilized to grow *P. aeruginosa* (LB) ^96^ and mimic the CF environment (SCFM) ^30^ and a media similar to the one provided to the mucosal tissue (DMEM) ^97^ (using ThermoFisher formulation with higher glucose, #11965). We also used minimal media, with glucose, N-acetyl-glucosamine, lactate, or single amino acids as single carbon sources, as well as glucose or N-acetyl-glucosamine supplemented with single amino acids. We further examined if other mucosal glycan monomers can serve as single carbon sources (i.e., mannose, xylose, fucose, galactose, N-acetylmannosamine, N-acetylgalactosamine and N-acetylneuraminate) ^98^. However, the model does not predict growth on any of these carbon sources.

To compare the results of the *in silico* gene essentiality analysis (i.e., mutant fitness compared to WT in each media condition) with the *in vitro* dataset (i.e., mutant fitness on tissue compared to mutant in LB), we transformed the *in vitro* data as follows. We used the 1.2-fold change as the threshold to consider the mutant phenotype on tissue to be different from the phenotype on LB. If the measured fold change was greater than the threshold, we considered the gene to be redundant in the tissue, while if the measured fold change was less than the inverse of the threshold, we considered the gene deletion to be growth-limiting in the tissue. Otherwise, we considered the mutant phenotype to be the same as predicted by the model for growth on LB media. To examine the sensitivity of the results to the threshold value, we repeated the analysis for 1.3- and 1.5-fold changes.

For the dimensionality reduction, we used the t-SNE function implemented in MATLAB with a Perplexity equal to 20 and LearnRate 100, and otherwise default parameters. To cluster the data, we used the k-means function implemented in MATLAB and reported the average cluster assignment after 1000 iterations. The optimal number of clusters was defined through silhouette analysis. All simulations for this study were performed on Mac Pro 32 GB in MATLAB 2017a and IBM ILOG Cplex 12.7.1 as a solver.

### Competition assays for image analysis, quantification of fitness, and cluster sizes

The competition assays used to quantify fitness and the distribution of cluster sizes (Fig. 3B-C; Fig. 4; Fig. S2) were performed as follows. A WT strain expressing mScarlet and competing strains expressing mNeonGreen (WT, Δ*wspF*, Δ*bifA*, Δ*wspF*Δ*pel*, Δ*wspF*Δ*psl*, and Δ*wspF*Δ*pel*Δ*psl*) were grown overnight in LB broth (containing gentamycin 30 µg/mL) from an LB plate, then cells were diluted 1:100 (WT, Δ*bifA*, Δ*wspF*Δ*pel*Δ*psl*) or 1:50 (Δ*wspF*, Δ*wspF*Δ*pel*, Δ*wspF*Δ*psl*) in new LB cultures and grown for around 3 h. Next, cells were spun down, washed in PBS buffer, and resuspended in PBS at an OD600 of 0.3. WT mScarlet was mixed with each competing strain in a 1:1 ratio used as inoculum for infections. Then, for each competition, three distinct Transwell inserts were infected using 1 µL of the 1:1 inoculum mix, and the infection progressed for 5 h.

Each Transwell insert used for infections contained tissue from CF HBE cells under differentiation for around 2-3.5 months. Each Transwell was incubated overnight (basal and apical) with P-ALI medium containing dye CellMask Deep Red (Life Technologies) at a concentration of 5 µg/mL for plasma membrane staining. In the morning, around 4 h before infection, the media with the dye was aspirated (both basal and apical), replaced with fresh P-ALI medium on the basal side, and left at ALI on the apical side. This process also removes the excess mucus.

After the infection ended, each Transwell insert was transferred to a glass-bottom 12- well black plate (1.5 coverslip) (IBL), where it was imaged using a Nikon Eclipse Ti2–E inverted microscope coupled with a Yokogawa CSU W2 confocal spinning disk unit. The microscope has a Prime 95B sCMOS camera (Photometrics). Four to six z-stacks were taken for each insert, with data collection on the GFP (mNeonGreen), Cy3 (mScarlet), and Cy5 (CellMask Deep Red) channels. Images were taken with a 20x water immersion objective with an N.A. of 0.95.

#### Image analysis

After data collection, initial processing involved manual inspection of stacks for image quality and selection of z-range where bacterial cells are present. Maximum intensity projections were then created for each field of view using Fiji v.2.0 ^99^. Projections were only generated for the two channels containing signals derived from the bacterial strains (mNeonGreen and mScarlet).

Next, we created a custom-made analysis pipeline to quantify the total area and cluster size for WT and competing strains within each competition comparison. The pipeline followed these steps. First, the z-projects were normalized with a mean-std normalizer before they were denoised, using the unsupervised denoising algorithm “noise2void” ^100^. The resulting model was saved and reused for all competition assays. Then, the denoised data was used for training the pixel classifier to detect background, cells, and blurry regions. The pixel classifier of choice was the Random Forest Classifier (RFC) ^101^ from scikit-learn ^90,102^. The training input for the model was the denoised data, the blurred version of the denoised data (Gaussian filter, sigma=2, scikit-image implementation), and the edge-enhanced version (Sobel filter, scikit-image implementation). The training annotation was generated manually. The training was then performed using GridSearch (scikit-learn implementation) to find the best parameters for the depth and the number of trees in the Random Forest. The RFC was trained for the two channels separately. The corresponding models were saved and used for all further predictions in the corresponding channels. The output cluster size was measured, and objects smaller than a small cell size were considered as artifacts and removed.

To quantify the area of the objects, we used the scikit-image implementation of connected component labeling (skimage.measure.label) ^103,104^, followed by extracting the area of the labels using region properties (skimage.measure.regionprops) ^105,106^. The single cluster areas were measured and saved for every field of view of every competition. This information was then used for figures containing single cluster size information. Specifically, all clusters for WT or competing strain within each competition were pooled together, and their cumulative distributions are shown in the figures (Fig. 4B, G; Fig. S2C). The data containing the cluster sizes was also used for calculating the percentage of small clusters (defined as clusters < 20 µm^2^) attributed to each strain within each of these competitions (Fig. 4C, E; Fig. S2D). Here, the total number of such clusters for WT and competing strain was counted, and the percentage of these attributed to each strain was calculated. This was done individually for each replicate (i.e., Transwell insert) within each competition assay, with values for WT and competing strain adding up to 100%.

Finally, the pipeline also includes calculating the total area covered by each strain within each field of view. For each field of view, the entire area covered by the competing strain was divided by the area covered by the WT to obtain the fitness ratios. The log2 transformation of these ratios is plotted in Fig. 3D, Fig. 4D, and Fig. S2A).

### Imaging bacteria growing mucus

Imaging bacteria growing on mucus (Fig. 1A) was done as follows. *P. aeruginosa* WT strain expressing mScarlet was grown in LB broth (containing gentamycin 30 µg/mL) from an LB plate for ∼four hours until reaching a dense culture, then cells were diluted to an OD600 of 0.25 and grown to deep stationary phase for 11 h. Next, cells were spun down, washed in PBS buffer, and resuspended in PBS at an OD600 of 1, which was used as inoculum for infections (same density as the one used for the Tn-seq infections). 1 µL of the inoculum was used, and infections progressed for three hours. The Transwell inserts used for infections contained tissue from CF HBE cells under differentiation for around 2.5 months. Each Transwell was incubated overnight (basal only) with P-ALI medium containing dye CellMask Deep Red (Life Technologies) at a concentration of 5 µg/mL for plasma membrane staining. Around one hour before starting the infections, the apical side of Transwells was stained with jacalin conjugated to fluorescein (Vector Laboratories) to label the mucus ^26^. The jacalin solution in PBS had a concentration of 50 µg/mL; incubation proceeded for five minutes, followed by complete aspiration. Transwells were then incubated at the ALI until the infections started. After the infection ended, Transwell inserts were imaged as described above, but using a 40x water immersion objective with N.A. of 1.15 with data collection on the GFP (fluorescein), Cy3 (mScarlet), and Cy5 (CellMask Deep Red) channels. We used Imaris (Bitplane) for 3D rendering of the z-stacks captured and Fiji for the display of images.

### Competition assays in AirGels without antibiotics

Competition assays in the absence of antibiotics in AirGels (Fig. 3E-F, Movie S1) were performed as follows. AirGels were stained overnight with the plasma membrane dye CellMask Deep Red (Life Technologies) diluted to a concentration of 5 µg/mL in the P- organoid medium (the medium used for AirGels differentiation and maintenance) both on the lumen and basal side. In the morning, around four hours before infections, the medium with the dye was aspirated, the basal side was replaced with fresh P-organoid medium, and the lumen was kept at the ALI until the infection started. Competition assays in the absence of antibiotics in AirGels used WT mScarlet and Δ*wspF* mNeonGreen. The inoculum was prepared as described above (OD600 = 0.3 of a 1:1 mix) and directly loaded into the lumen of AirGels (using 0.5 µL) like we previously described ^26^. This small volume kept the lumen at the ALI. This inoculum led to a multiplicity of infection of approximately 1. The infection proceeded for five hours at 37 °C with 5% CO2 in the cell culture incubator. As we noticed before ^26^, condensation appears on the PDMS chip during imaging. Therefore, to prevent dripping water from disrupting the ALI conditions inside the lumen, we placed pieces of Kimtech Science Kimwipes (Kimberly–Clark Professional) in the inlet ports of AirGels after infections. After the initial five hours of infection, AirGels were moved to the microscope for imaging. This was done with a UNO-T-H-CO2 stage-top incubator (Okolab), maintaining the same incubation conditions (37°C with 5% CO2 and humification). Chips were imaged for at least eight hours, with the acquisition for each position covering a z-range of 50-60 µm every 15 minutes. Three AirGels were imaged (one position each) with our 20x water immersion objective (N.A. of 1.15) and data collection on the GFP (mNeonGreen), Cy3 (mScarlet), and Cy5 (CellMask Deep Red) channels. A custom-made water injector kept the water immersion on the objective throughout the acquisition.

The data processing was done using Elements v.5.21.02 (Nikon Instruments) and Fiji. Briefly, maximum intensity and average intensity projections for each stack were created using Nikon Elements software. Maximum intensity projections were used for visualization (Movie S1), while average intensity projections were used for quantification (Fig. 3F), both done in Fiji. Fluorescence was quantified for each time point in each projection using the “Mean gray value” function (Fig. 3F-left). Only the mNeonGreen and mScarlet channels were used for this quantification. Then, within each field of view and time point, Δ*wspF* fluorescence was divided by WT fluorescence; this ratio was log2- transformed and used for plotting (Fig. 3F-right)

### Growth and cytotoxicity of P. aeruginosa strains in AirGels

Assays measuring growth and cytotoxicity of individual strains in AirGels (Fig. S3, Movies S2-S4) were performed as follows. AirGels were stained overnight with the plasma membrane dye as described above. In the morning, around four hours before infections, the medium with the membrane dye was aspirated and replaced (both luman and basal) with the P-organoid medium containing propidium iodide (PI) at a concentration of 2.5 µg/mL. This staining proceeded until ∼one hour before the infection, when the medium was again aspirated, the basal side was replaced with fresh P-organoid medium, and the lumen was kept at the ALI. The infections used WT and Δ*wspF* strains expressing mNeonGreen. The inoculum was prepared as described above (OD600 = 0.3 for each, infections done separately) and directly loaded into the lumen of AirGels (using 0.5 µL). The infection proceeded for three hours at 37 °C with 5% CO2 in the cell culture incubator. After the initial three hours of infection, AirGels were moved to the microscope, and chips were imaged for at least eight hours (using a stage-top incubator), with the acquisition for each position covering a z-range of 50-60 µm every 15 minutes. Two AirGels were imaged (one position each) with our 20x water immersion objective (N.A. of 1.15) and data collection on the GFP (mNeonGreen), Cy3 (PI), and Cy5 (CellMask Deep Red) channels. Maximum intensity projections for each stack were created using Nikon Elements software, and projections were further processed in Fiji.

### Antibiotic tolerance in AirGels

Antibiotic tolerance in AirGels was performed as follows. AirGels were prepared for infections as described above. Then, two types of infection experiments were performed: (i) single-strain infections using WT or Δ*bifA*, both expressing mNeonGreen (Fig. 5D-F; Movie S5); or (ii) co-infections with both WT and Δ*wspF* together, expressing mScarlet and mNeonGreen, respectively (Fig. 6; Movie S6-S7). The inoculum preparation and loading followed what was described above (using 0.5 µL of OD600 = 0.3, either single strains or mixed 1:1). The infection proceeded for five hours at 37 °C with 5% CO2 in the cell culture incubator, after which the AirGels were moved to the microscope for imaging, ciprofloxacin was added to the basal side of the chips (at a final concentration of 1 µg/mL mixed into the P-organoid medium). AirGels were imaged for least 15 hours (using stage-top incubator), with the acquisition for each position covering a z-range of 50-60 µm every 15 minutes. Six AirGels were imaged (one to four positions per chip) for each treatment in the single strain infections, and two AirGels were imaged (four positions per chip) for the WT and Δ*wspF* co-infections. To avoid basal medium leakage inside the lumen and consequent disruption on the ALI throughout the extended length of the experiment, we always used data from AirGels that had not been severely damaged by the time the antibiotic was added. Imaging used our 20x water immersion objective (N.A. of 1.15) and data collection on the GFP (mNeonGreen), Cy3 (mScarlet, only for co-infections), and Cy5 (CellMask Deep Red) channels.

3D rendering of representative z-stacks was done with Imaris (Bitplane) using the same thresholding for compared regions (Fig. 6B, Movie S6). The data not related to 3D rendering was processed using Elements (Nikon Instruments) and Fiji. The maximum intensity and average intensity projections for each stack were created using Elements.

As for the growth experiments in AirGels, maximum intensity projections were generally used for visualizations, while average intensity projections were used for quantifications, both done in Fiji. First, fluorescence was quantified for each time point in each projection using the “Mean gray value” function. Only the mNeonGreen channel was used for the quantification of single-strain infections, while both mNeonGreen and mScarlet channels were used for quantification of co-infections. Fluorescence patterns always followed an immediate increase for around four hours (time required for the ciprofloxacin to diffuse through the hydrogel, reach, and start acting on the cells), followed by a decrease in fluorescence at distinct rates depending on the strain/conditions. We then performed min-max normalization on each curve derived from each position imaged. For single-strain infections, distinct positions within the same replicate were combined by using their mean min-max normalized fluorescence, and the six replicates were used for plotting (Fig. 5F). For co-infections, given our interest in the Δ*wpsF*-mediated tolerance heterogeneity within the same AirGel, each position was plotted separately (Fig. 6E). Finally, we extracted the time at which the normalized fluorescence values of the WT dropped to 50% or less from its maximum for distinct conditions (which we called *τ*, Fig. 6F). These included data from WT (expressing mNeonGreen) exposed to ciprofloxacin in single-strain infections and WT (expressing mScarlet) exposed to the antibiotic in co-infections with Δ*wspF*. For the co-infections, we split the data into positions where Δ*wspF* formed small or large biofilms.

### Data processing and plotting

Data processing was generally performed in Python (version 3.9) ^91^, using a combination of the Pandas (version 1.4.4) ^89^, NumPy (version 1.21.5) ^107^, SciPy (version 1.7.3) ^108^, and scikit-learn (version 1.3.2) ^90^ libraries. In addition, Microsoft Excel (version 16.39) was also used.

All the plots displayed in the manuscript were prepared using Matplotlib (version 3.5.2)^109^ and Seaborn (version 0.12.0) ^110^. Plot legends and their display organization within each figure were adjusted using Adobe Illustrator (Adobe). The same software was used to draw all the illustrations shown in the manuscript.

### Statistical analysis

All statistical analyses were performed using Python. Either Welch unpaired t-tests or 1-way analysis of variance (ANOVA) with post hoc Tukey honestly significant difference (HSD) test for multiple comparisons were used, as specified in the figures’ legends.

## Acknowledgments

We thank Zaïnebe Al-Mayyah for laboratory technical assistance, the Lausanne Genomic Technologies Facility (GTF) for sequencing, and members of the Persat lab for constructive feedback throughout the development of the project. We also thank Dianne K. Newman, Megan Bergkessel and Carey Nadell for comments on the manuscript.

## Author contributions

Conceptualization: LAM, AP

Data curation: LAM, EV, ES

Formal Analysis: LAM, EV, ES

Funding acquisition: VH, AP

Investigation: LAM, EV

Methodology: LAM, EV, AD, ES, TR, TD

Project administration: LAM, AP

Supervision: VH, AP

Visualization: LAM, EV, ES

Writing – original draft: LAM, AP

Writing – review & editing: LAM, EV, VH, AP

## Declaration of interests

Authors declare that they have no competing interests.

## Funding

Swiss National Science Foundation (SNSF): 310030_189084 (AP) NCCR AntiResist (AP)

European Molecular Biology Organization (EMBO) Postdoctoral Fellowship ALTF 12- 2022 (LAM)

Swiss National Science Foundation (SNSF): 200021_188623 (VH) NCCR Microbiomes (VH)

## Notes

### Competing Interest Statement

The authors have declared no competing interest.

