## Supplementary material for "*Pseudomonas aeruginosa* faces a fitness trade-off between mucosal colonization and antibiotic tolerance during airway infections"

**Supplementary figures**

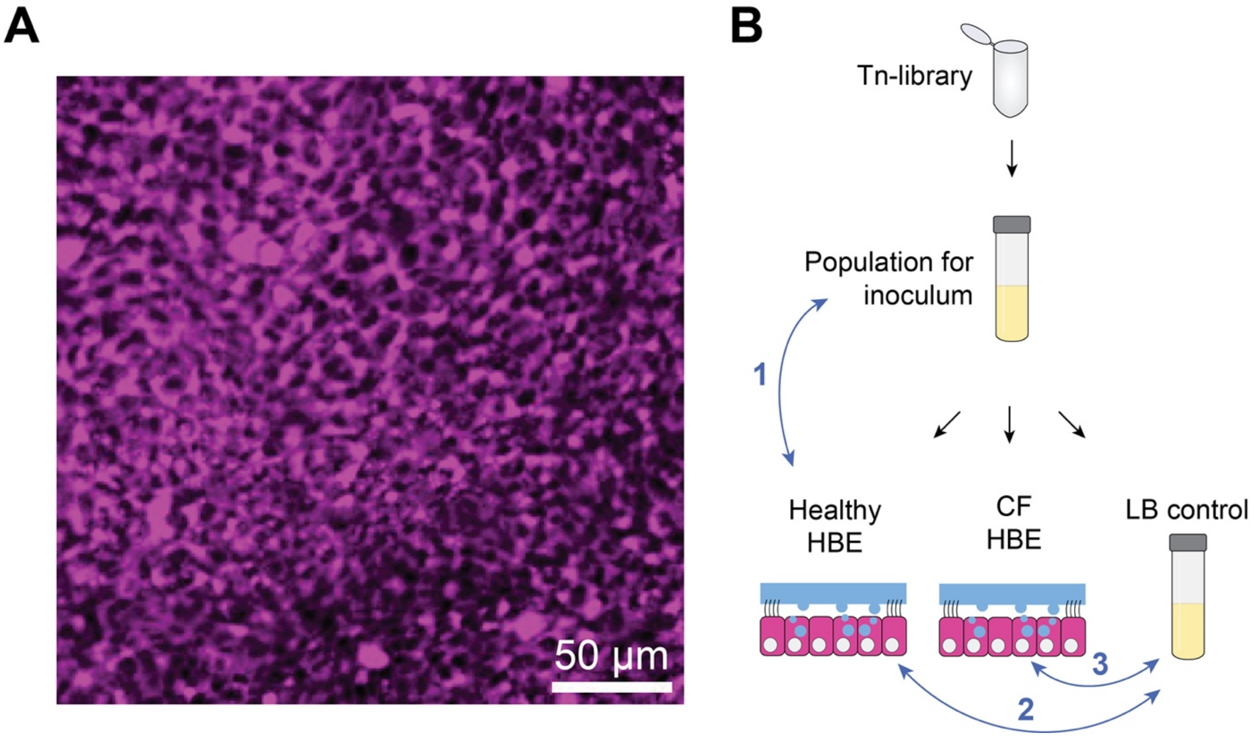


**Figure S1. Detailed procedure for the Tn-seq during mucosal colonization. A.** Representative confocal image of (z-slice) showing intact epithelium after 11 h of infection with the Tn-library. **B.** Exhaustive illustration of experiments and control conditions for the Tn-seq during mucosal colonization. All five samples were sequenced (Tn-library, population used for inoculum, healthy HBE, CF HBE, and LB control). Blue arrows represent the comparisons (1-3) made using the TRANSIT software to assess the conditional essentiality of genes. In comparison 1, the inoculum was used as the control; in comparison 2 and 3, the "LB control" condition was used as the control.

Abbreviations: CF, cystic fibrosis; HBE cells, human bronchial epithelial cells.

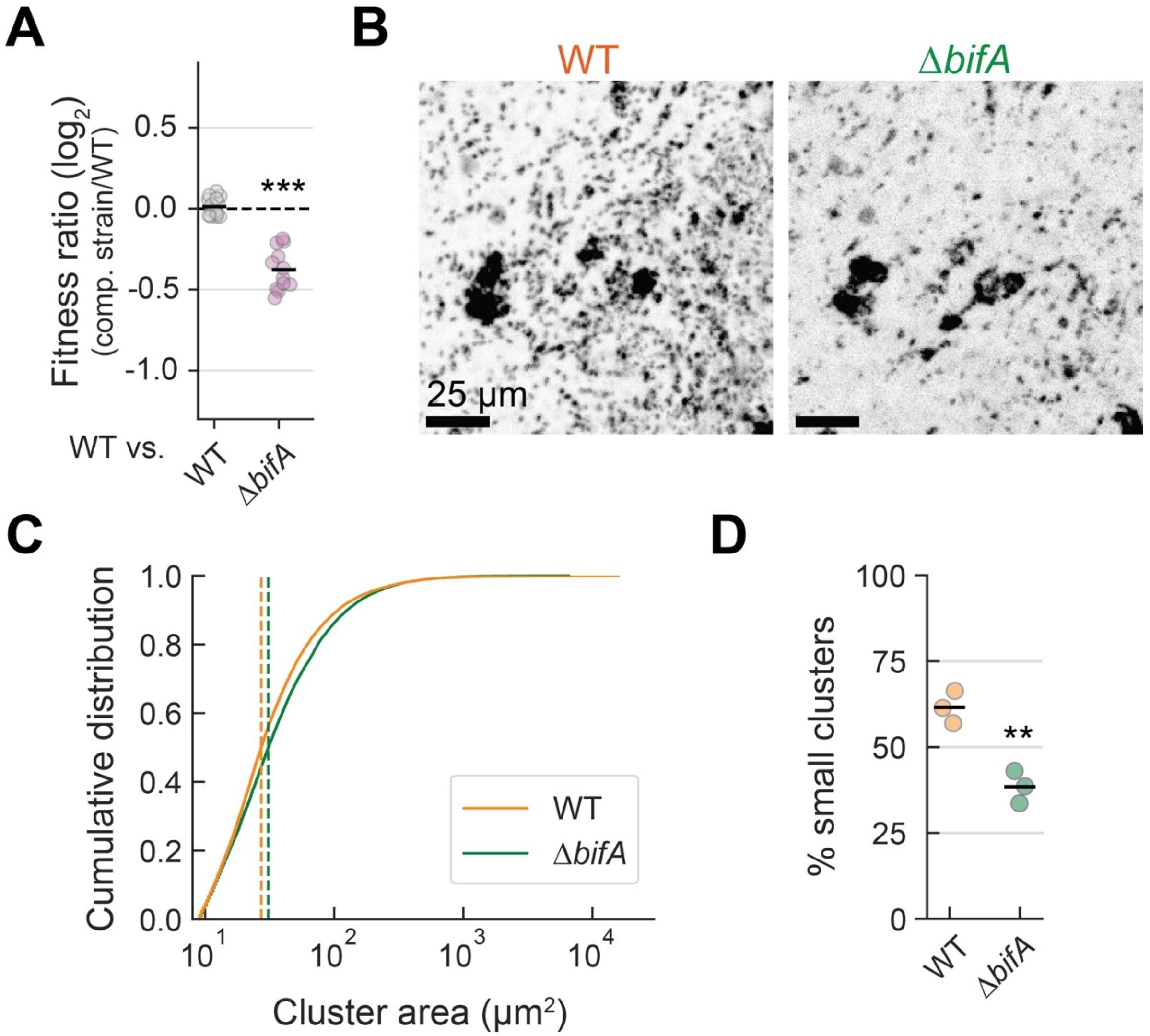


**Figure S2. Biofilm fitness quantification during mucosal colonization. A.** Fitness ratio based on the quantification of the total area measured from surface coverage by Δ*bifA*-mNeonGreen relative to WT-Scarlet during competition assays at the mucosal surface (HBE cultures). Each data point represents one imaged field of view (n = 15 for WT vs. WT; *n* = 12 for WT vs. Δ*bifA*) distributed within three biological replicates. The data for the WT-mNeonGreen vs. WT-mScarlet competition is the same from Fig. 3E but is shown here for comparison. Horizontal black lines mark the mean fitness ratio for each condition. **B.** Representative images of WT and Δ*bifA* competition assays in HBE cultures (maximum intensity projection; same field of view). **C.** Cumulative distributions for cluster size formed by WT and Δ*bifA*. Distributions include three biological replicates. Vertical lines represent the median cluster-size for each strain. **D.** Percentage of small clusters (smaller than 20 μm$^2$) for WT and Δ*bifA* populations. Each data point represents an independent biological replicate ($n$ = 3); horizontal black lines mark their mean. Statistics: panel A and D, Welch unpaired t-test (*** $p < 0.001$, ** $p < 0.01$).

Abbreviations: HBE cells, human bronchial epithelial cells; WT, wild-type.

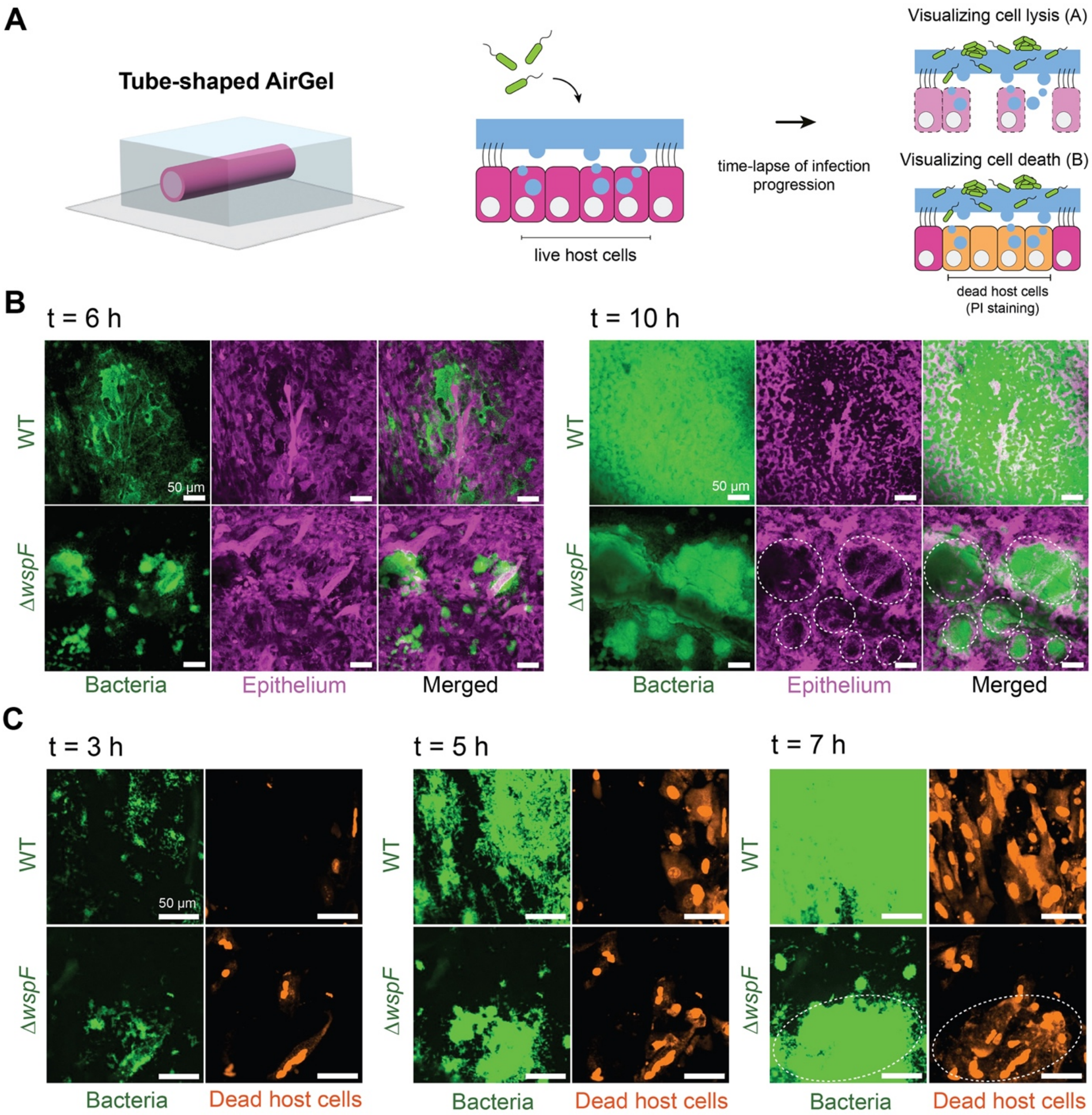


**Figure S3. Biofilms mechanically damage epithelia while constraining pathogenicity. A.** Experimental design for imaging epithelial damage upon WT and Δ*wspF* infections in AirGels. AirGels were infected WT or Δ*wspF* expressing mNeonGreen, and infections were monitored for 11 hours for the observation of lysis and cell viability (propidium iodide staining). **B.** Representative maximum intensity projection images showing differences in growth and subsequent epithelial cell lysis by WT (top) and Δ*wspF* (bottom). Note that after 10 h of infection, WT had completely taken over the surface and had lysed the epithelial cells. In contrast, Δ*wspF* formed large expanding biofilms that opened up nodules that stretched out within the tissue (white circles). **C.** Representative maximum intensity projection images showing differences in growth and the killing of epithelial cells by WT (top) and Δ*wspF* (bottom). Note the uniform cytotoxic effect to the WT due to its faster spreading, while Δ*wspF* biofilms showed only cytotoxic activity towards epithelial cells in the immediate vicinity of the nodules (white circles). Abbreviations: PI, propidium iodide; WT, wild-type.

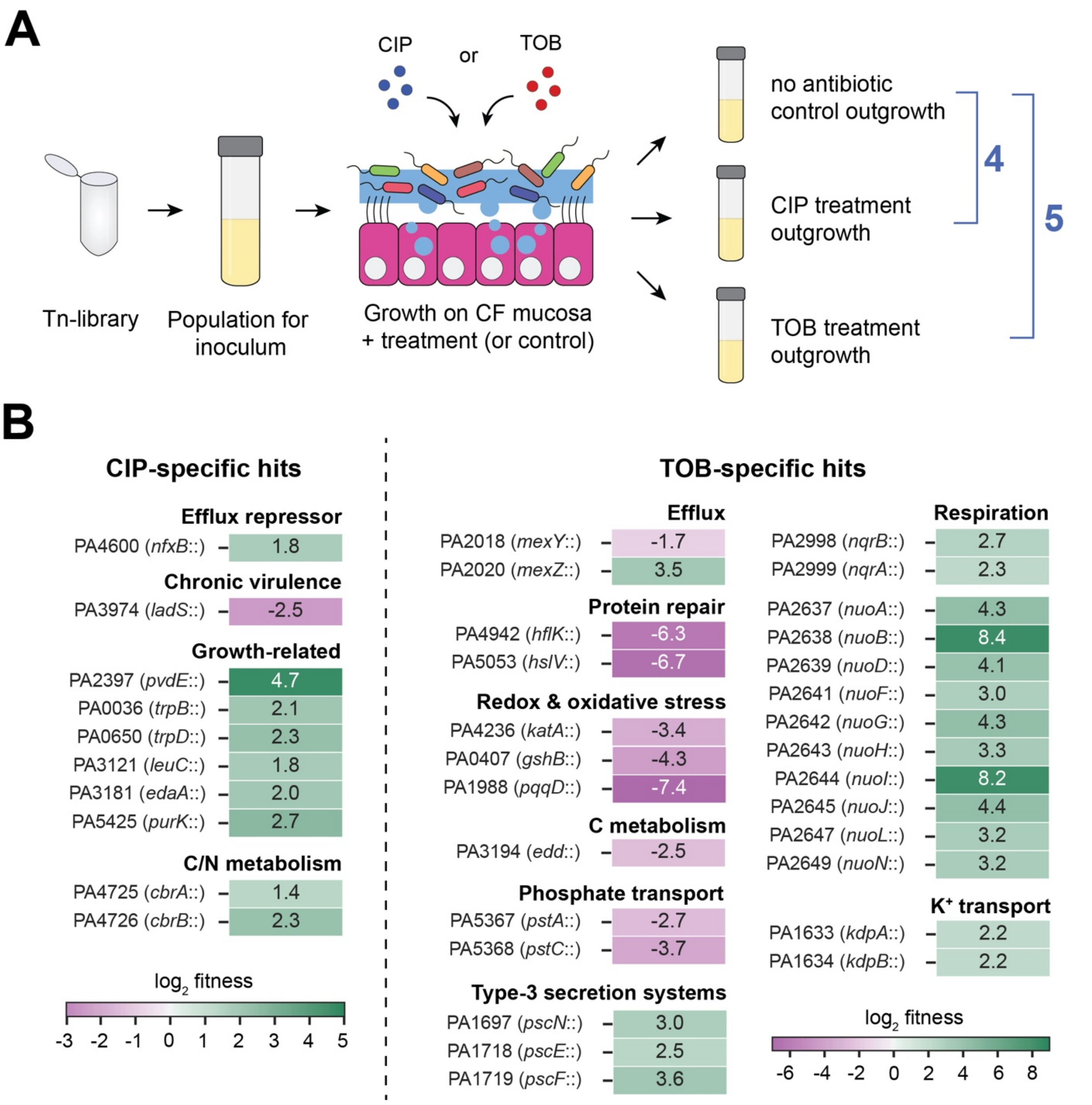


**Figure S4. Tn-seq analysis identifies regulators of antibiotic adaptation at the mucosal surface. A.** Full experimental design of the Tn-seq during antibiotic tolerance at the mucosal surface. The library was grown on CF HBE cells, treated with CIP or TOB (or no treatment for the control), and then an outgrowth step on LB was performed for all samples. The three outgrowth samples were sequenced. Blue lines represent the comparisons made using the TRANSIT software to assess the conditional essentiality of genes (comparisons 4-5), and the “no antibiotic” condition was used as the control. **B.** Fitness effects of transposon insertions in representative genes and their categories discovered in our antibiotic tolerance Tn-seq. Genes are separated by CIP and TOB-specific hits. These do not represent all the genes that made the significance cutoff. See Table S4 for the complete dataset.

Abbreviations: CIP, ciprofloxacin; CF, cystic fibrosis; HBE cells, human bronchial epithelial cells; TOB, tobramycin.

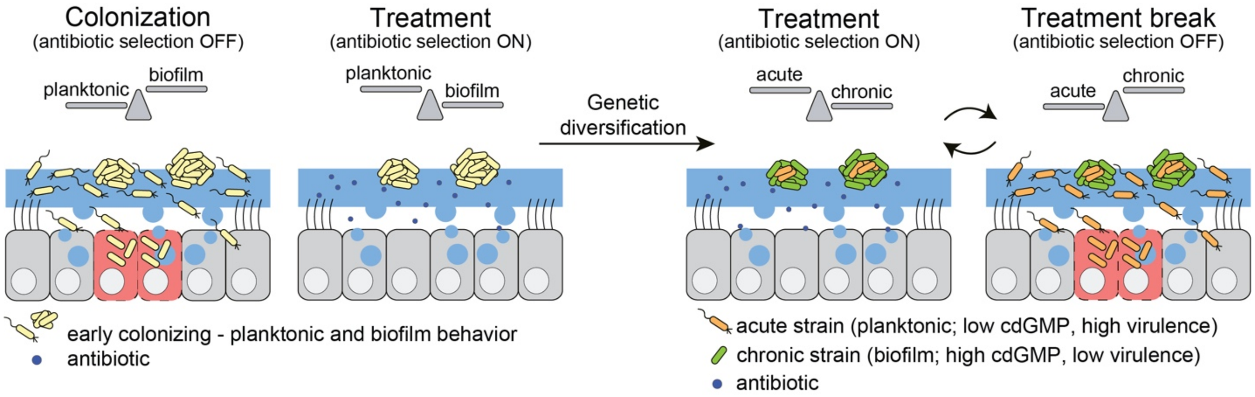


**Figure S5. Model of the colonization-tolerance trade-offs for biofilms and its consequences for genetic diversification during chronic infections. Left.** During colonization, planktonic behavior is beneficial as it allows better spread on the mucosa, eventually leading to the killing of epithelial cells. However, upon antibiotic treatment selection, biofilm lifestyle, which minimizes tissue damage, is selected. **Right.** Long-term infection leads to genetic diversification, where strains displaying different genotypes co-exist. Such genotypes display phenotypic lifestyles (acute or chronic behavior). During antibiotic treatment selection, biofilm-forming strains may protect genotypes associated with acute behavior. We hypothesize that upon removal of the antibiotic selection (either by treatment break and/or mutation leading to resistance), acute behavior is again beneficial, potentially leading to lung exacerbations.

**Additional supplementary files**

**Supplementary tables**

**Table S1.** Complete list of genes identified in the Tn-seq of *P. aeruginosa* growing at the mucosal surface (healthy) compared to the liquid culture inoculum reference.

**Table S2.** Complete list of genes identified in the Tn-seq of *P. aeruginosa* growing on CF and non-CF HBE cultures compared to an LB control culture.

**Table S3.** Full dataset for the comparison between *in silico* fitness simulations and fitness found in the Tn-seq experiment.

**Table S4.** Complete list of genes identified in the Tn-seq measuring the tolerance of *P. aeruginosa* to ciprofloxacin or tobramycin in comparison to the untreated control.

**Table S5.** Strains, plasmids, and primers used in this study.

**Supplementary movies**

**Movie S1.** Growth of WT (orange) and Δ*wspF* (green) during AirGel co-infections.

**Movie S2.** Growth of WT (top) and Δ*wspF* (bottom), both in green, during separate AirGel infections.

**Movie S3.** Visualization of tissue lysis by WT (top) and Δ*wspF* (bottom).

**Movie S4.** Epithelium cell death measured with propidium iodide over the course of AirGel infections by WT (top) or Δ*wspF* (bottom).

**Movie S5.** Representative movies displaying the tolerance levels WT (top) and Δ*bifA* (bottom) upon ciprofloxacin treatment in AirGels. Both strains are shown in green.

**Movie S6.** Representative 3D rendering movies showing differences in tolerance to ciprofloxacin of mixed infections forming large (left) or small (right) biofilms. The two distinct regions shown were collected in the same AirGel, located millimeters apart. WT is shown in orange and Δ*wspF* is shown in green, and the epithelium is shown in magenta.

**Movie S7.** Representative maximum intensity projection movies showing differences in tolerance to CIP of mixed infections forming large (top) or small (bottom) biofilms. The two distinct regions shown were collected in the same AirGel, located millimeters apart. WT is shown in orange and Δ*wspF* is shown in green.
